# Differing effects of parasite-parasite interaction types on the spatial epidemiology of co-circulating parasites

**DOI:** 10.64898/2026.04.02.716128

**Authors:** Giacomo Zilio, Javier Zabalegui Bayona, Laure Rousseau, Jérôme Raichle, Claire Gougat-Barbera, Alison B. Duncan, Andrew D. Dean, Oliver Kaltz, Andy Fenton

## Abstract

Interactions among co-circulating parasite species influence infection risk and disease progression. Interactions can occur within hosts, e.g. altering susceptibility, or indirectly through host demography or movement, potentially affecting landscape-scale transmission. Despite their ubiquity, the spatial implications of these interactions have received limited attention. We combine spatially-explicit modelling with laboratory experiments to investigate how parasite–parasite interactions influence disease spread. We model within-host, demographic, and dispersal-related interactions across a linear landscape, showing that within-host interactions modifying host susceptibility have the strongest effects on parasite prevalence, spatial heterogeneity, and rate of spread. These effects are amplified when parasites invade sequentially, generating pronounced patch-level priority effects. We tested these predictions experimentally using a protist host (*Paramecium caudatum*) and two bacterial parasites (*Holospora undulata* and *H. obtusa*). Consistent with model predictions, we found that *H. obtusa* reduces the overall prevalence and spread of H. undulata across a landscape of interconnected patches, likely through reductions in host susceptibility, and evidence of priority effects, observing reduced *H. undulata* prevalence when introduced after *H. obtusa*. Our results highlight that parasite-parasite interactions can have important implications for the landscape-scale epidemiology of co-circulating parasites, but the effect magnitude depends on the interaction type and the timing of invasion.

## INTRODUCTION

It is well known that co-circulating parasite species can interact within individual hosts to alter the probability of infection or the severity of disease progression, with potential implications for parasite transmission [1 -5]. However, how interactions between parasites scale up to influence the landscape-level spread of disease is largely unknown.

Intuitively, it makes sense that parasite-parasite interactions could impact disease spread over larger scales. As outlined by Daversa et al [6], there are a number of key epidemiologically-relevant processes that influence the spatial spread of infectious diseases, and there is now a large body of evidence showing how the presence of co-circulating and co-infecting parasites can influence each of these processes (Figure 1; [5]). These effects can arise through direct interactions at the within-host level (e. g., one parasite alters host susceptibility to infection by other parasites [7-10]). However there may also be indirect, demographic interactions acting at the host population level, whereby one parasite alters the survival or abundance of hosts available for infection by other parasites [11-13] or alters the propensity for host movement [14-17]. Hence, co-circulating parasites can interact even without simultaneous infection of the same host individual [5]. Given the ubiquity of co-circulating parasites, and the diverse ways they can interact, the cumulative effects of these processes could strongly alter the spatiotemporal dynamics of disease transmission across systems of conservation, economic and public health concern. However, there has been little attempt to consider the consequences of parasite-parasite interactions on the spatiotemporal dynamics of co-circulating parasite species.

**Figure 1.**
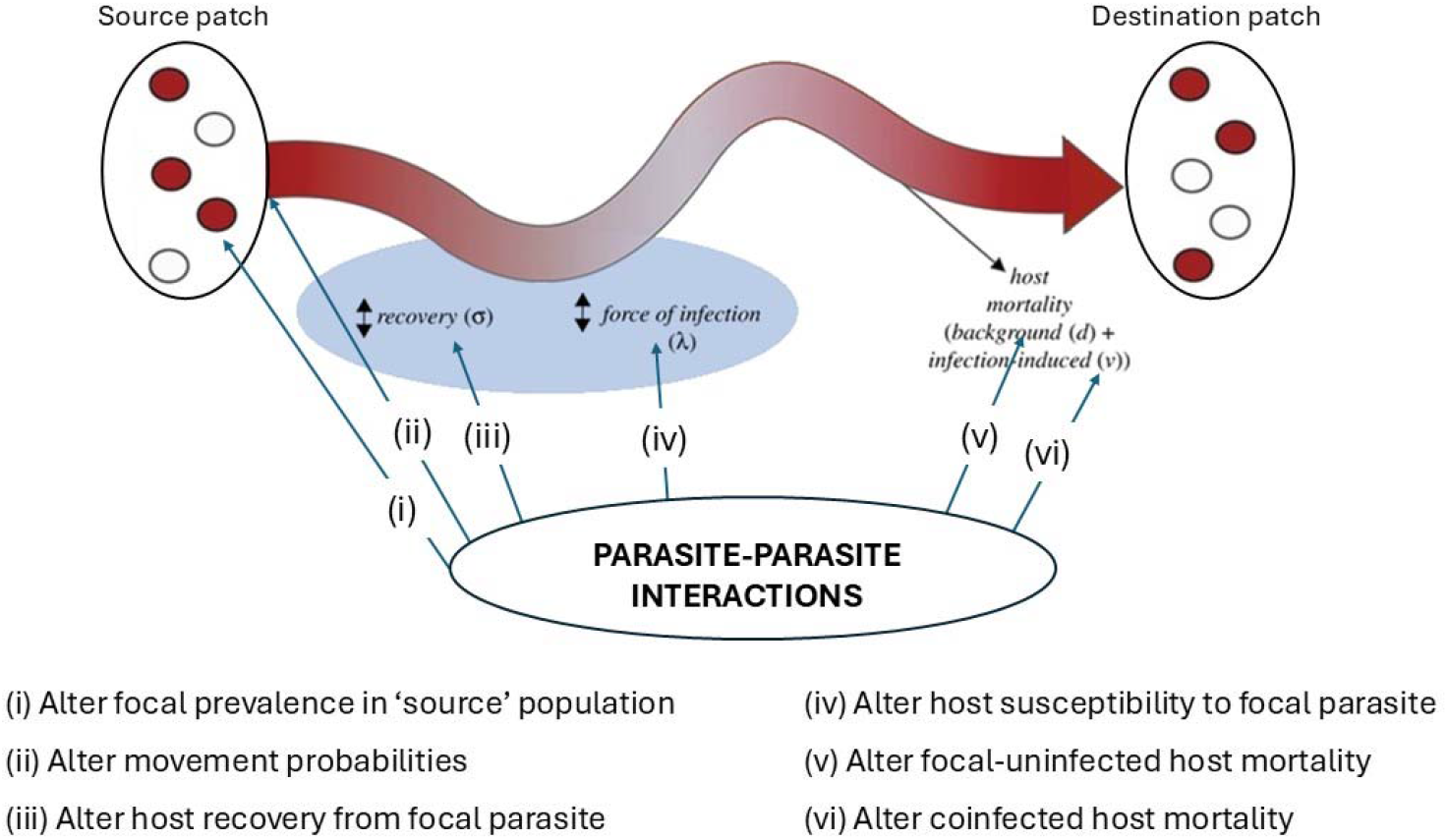
Conceptual framework showing different ways in which parasite-parasite interactions may affect the spatial spread of a parasite at different points of the dispersal pathway between source and destination patches. Modified from Daversa et al [6].

To date, theoretical studies exploring the spatial impact of parasite-parasite interactions have considered a relatively limited range of interaction types, and empirical work has been largely observational and not directed towards testing the theoretical predictions. In terms of general theory, Poletto et al [18,19] developed stochastic metapopulation models of two co-circulating parasite species interacting via protective cross-immunity (i.e., a ‘within-host interaction’ [5]), such that infection by one parasite prevented infection by the other parasite (no simultaneous co-infection of hosts by both species). They showed that low host migration rates favour parasite strains with longer infectious periods (and lower transmissibility per unit time, for the same overall R_0_), whereas high migration rates favour strains with shorter infectious periods and higher transmissibility [18]. Furthermore, these effects were moderated by the degree of cross-immunity [19]. Conversely, Menezes & Rangel [20] explored the effect of two co-circulating pathogens on the spatial dynamics of three host species undergoing rock-paper-scissors interactions, allowing for co-infection of each species. They showed that if co-infected hosts suffer increased mortality compared to single-infected hosts, this results in discrete spatial regions of hosts infected by one or other pathogen, separated by interfaces of co-infected hosts. To date though there has been little theory directly assessing how different forms of parasite-parasite interactions affect host and parasite spatial dynamics.

Empirically, previous studies have considered the large-scale spatial overlap of co-circulating infections, often from a public health perspective [21,22] (although see also [23]), seeking to identify hotspots of potential co-exposure or to draw inferences about potential interactive effects (e. g., spatial zones of apparent exclusion or enhanced co-occurrence, which may be indicative of antagonistic or synergistic interactions between the parasites). However, such population-level patterns can be hard to interpret mechanistically. Both theory [24-27] and empirical studies [28, 29] have shown that within-host co-infection interactions can generate counter-intuitive patterns in parasite occurrence at the host population level (see also [26]). At a local scale, Keegan et al [30] analysed spatially-explicit data of wood mice (Apodemus sylvaticus) and two co-circulating parasites, a nematode and a coccidian, where previous experiments [31] showed the nematode suppresses shedding of coccidial infective stages (oocysts) from co-infected hosts. They found that high local prevalence of the nematode was associated with reduced coccidial infection risk, but only within restricted ranges (up to ∼50m), approximating the home range size of the host. Hence, within-host suppression of oocyst shedding from co-infected hosts led to a local reduction of between-host coccidial transmission. How such local effects scale up to the landscape level remains an open question.

Clearly then, existing theoretical and empirical studies suggest that parasite interactions have the potential to alter parasite spatiotemporal dynamics, but the overall picture remains incomplete. Here we combine generalised spatially-explicit theory and experiments using a tractable multiparasite lab system, to assess how different forms of parasite-parasite interaction impact parasite spatial dynamics. Our model considered three types of interaction (within-host, demographic or dispersal-related) between two parasite species dispersing over a linear landscape of connected patches. We found parasite spatial dynamics were affected primarily by a within-host interaction, whereby one parasite alters host susceptibility to another, impacting its landscape-wide prevalence, spatial heterogeneity, and dispersal across the landscape. Furthermore, we found the outcomes depended on the epidemiological scenario, such that sequential arrival of the parasites magnified interaction effects, leading to strong priority effects with landscape-scale consequences.

We then evaluated the model predictions empirically, using a lab experimental system involving the freshwater protist *Paramecium caudatum*, with two co-circulating bacterial parasites (*Holospora* sp.), known to strongly interact with each other across scales [32]. Mirroring the setup of our model, we quantified the role of parasite-parasite interactions on the spread of infection across a linear landscape of interconnected microcosms. As predicted, we found that the presence of one parasite (*H. obtusa*) consistently limited local outbreaks and the rate of spread among patches of the other species (H. undulata). Together, these results show that parasite spatiotemporal dynamics can be altered by the presence of other co-circulating parasites, particularly if they interact directly at the individual host level, and that the magnitude of these effects depend on the timing of invasion of the parasites relative to each other.

## METHODS

### Epidemiological model

We developed a generalised stochastic metapopulation-style model, similar conceptually to that of Poletto et al [18, 19], by modelling the patch-level dynamics of two interacting parasites, allowing for simultaneous co-infection of individual hosts (Figure 2a), with host and parasite dispersal between patches. We considered the patches to be spatially arranged in a linear chain (Figure 2b), allowing us to directly quantify rates of parasite spread from an initial point of introduction under alternative parasite-parasite interaction scenarios, and enabling conceptual links with our experimental design (see below).

**Figure 2.**
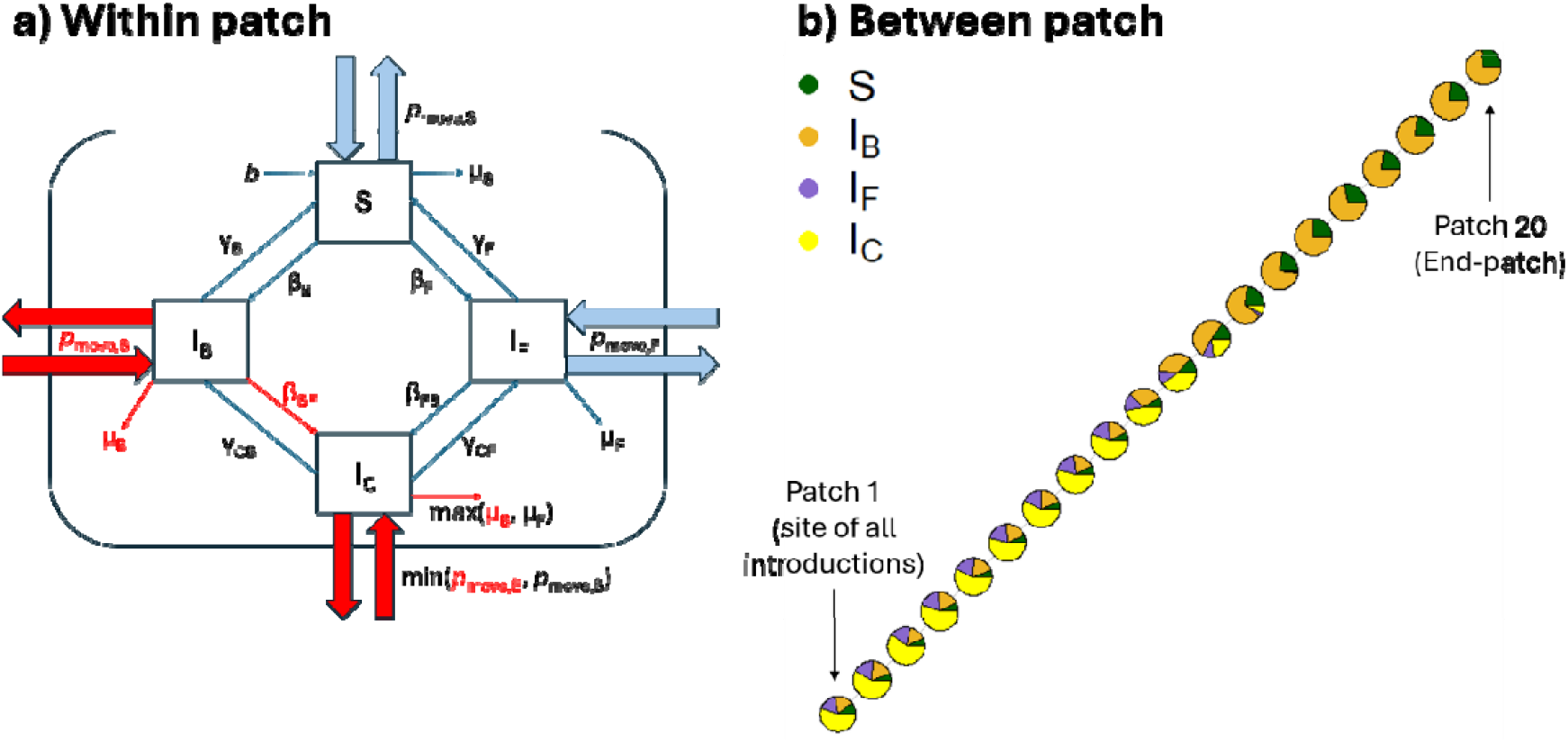
Schematics of (**a**) the within-patch epidemiological model, and **(b)** the between-patch network. For the within-patch model, the red arrows and labels show the co-infection and co-circulation interactions explored. For the between-patch figure, the pie charts show the relative proportions of the different host classes at each patch around the mid-point of invasion of the focal parasite (timestep 50 post introduction into Patch 1).

Our model (see Figure 2a for a model schematic, Table 1 for parameter definitions, and the Supplementary Information 1 for model details) tracked the dynamics of two co-circulating parasites: a focal parasite, ‘*F*’ (where *I*_*F*_ represents the number of hosts infected by the focal parasite only, in a given patch), whose epidemiological dynamics we are primarily interested in, and a background parasite, ‘*B*’ (where *I*_*B*_ represents the number of hosts infected by the background parasite only, in a given patch), which may alter the dynamics of the focal parasite under various interaction scenarios (see below). The within-patch dynamics of both parasites followed a general SIS (Susceptible-Infected-Susceptible) framework, with an additional host class,, representing individual hosts simultaneously co-infected with both parasites.

**Table 1.**
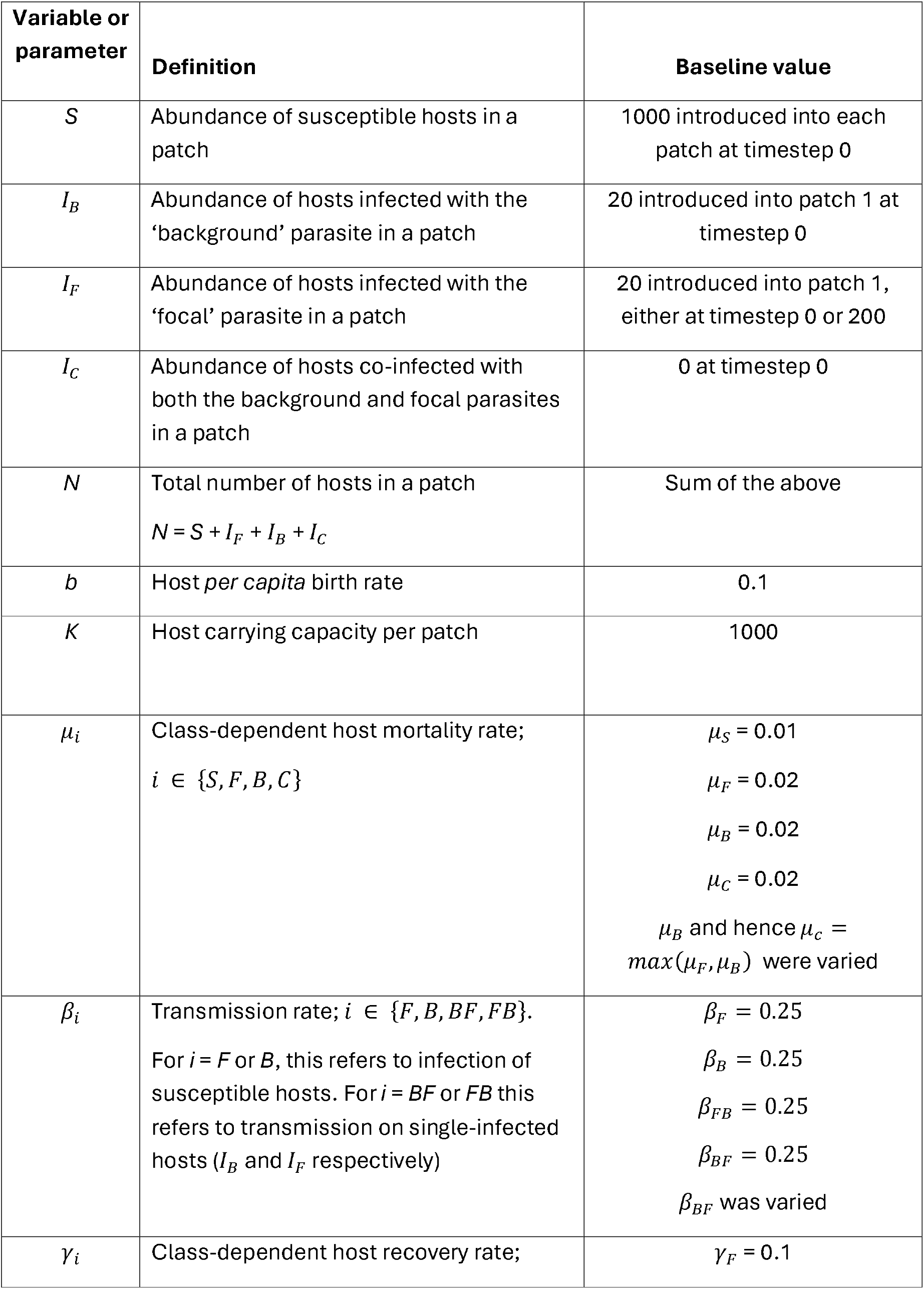
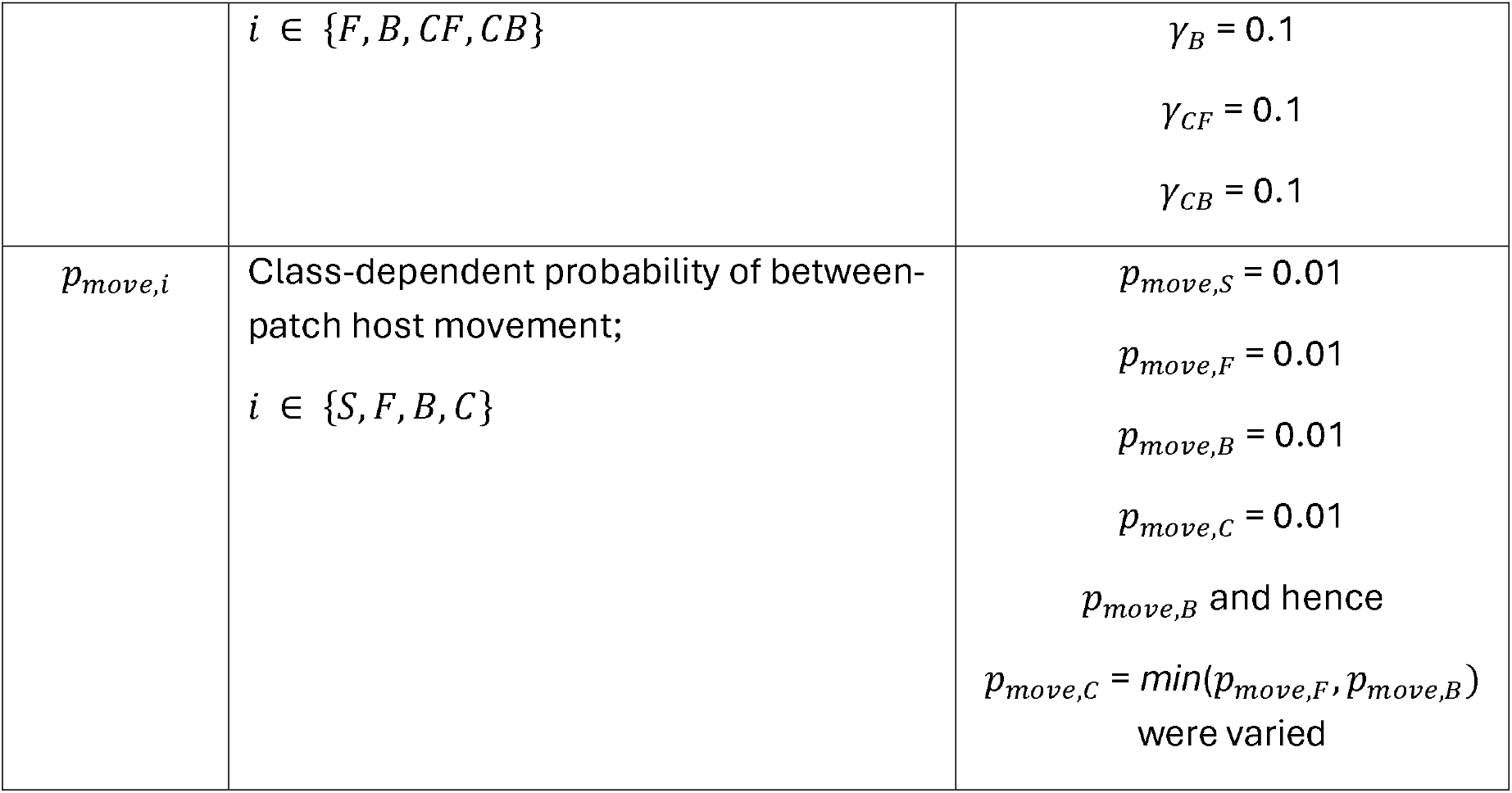
Variable and parameter definitions and baseline values used in the model simulations.

| Variable or parameter | Definition | Baseline value |
| --- | --- | --- |
| $S$ | Abundance of susceptible hosts in a patch | 1000 introduced into each patch at timestep 0 |
| $I_B$ | Abundance of hosts infected with the ‘background’ parasite in a patch | 20 introduced into patch 1 at timestep 0 |
| $I_F$ | Abundance of hosts infected with the ‘focal’ parasite in a patch | 20 introduced into patch 1, either at timestep 0 or 200 |
| $I_C$ | Abundance of hosts co-infected with both the background and focal parasites in a patch | 0 at timestep 0 |
| $N$ | Total number of hosts in a patch<br>$N = S + I_F + I_B + I_C$ | Sum of the above |
| $b$ | Host *per capita* birth rate | 0.1 |
| $K$ | Host carrying capacity per patch | 1000 |
| $\mu_i$ | Class-dependent host mortality rate;<br>$i \in \{S, F, B, C\}$ | $\mu_S = 0.01$<br>$\mu_F = 0.02$<br>$\mu_B = 0.02$<br>$\mu_C = 0.02$<br>$\mu_B$ and hence $\mu_C = \max(\mu_F, \mu_B)$ were varied |
| $\beta_i$ | Transmission rate; $i \in \{F, B, BF, FB\}$.<br>For $i = F$ or $B$, this refers to infection of susceptible hosts. For $i = BF$ or $FB$ this refers to transmission on single-infected hosts ($I_B$ and $I_F$ respectively) | $\beta_F = 0.25$<br>$\beta_B = 0.25$<br>$\beta_{FB} = 0.25$<br>$\beta_{BF} = 0.25$<br>$\beta_{BF}$ was varied |
| $\gamma_i$ | Class-dependent host recovery rate; | $\gamma_F = 0.1$ |
| | $i \in \{F, B, CF, CB\}$ | $\gamma_B = 0.1$<br>$\gamma_{CF} = 0.1$<br>$\gamma_{CB} = 0.1$ |
| $p_{move,i}$ | Class-dependent probability of between-patch host movement;<br>$i \in \{S, F, B, C\}$ | $p_{move,S} = 0.01$<br>$p_{move,F} = 0.01$<br>$p_{move,B} = 0.01$<br>$p_{move,C} = 0.01$<br>$p_{move,B}$ and hence<br>$p_{move,C} = \min(p_{move,F}, p_{move,B})$<br>were varied |

We assumed frequency-dependent transmission (as in Poletto et al [18, 19]), based on the numbers of susceptible, infected and total hosts within each patch at each time step. represents the baseline *per capita* transmission rates for parasite *i* on susceptible hosts, and represents the transmission rate of parasite i to hosts already infected by parasite *j*. Thus, differing from represents a within-host interaction such that prior infection by parasite *j* altered host susceptibility to parasite *i*. We note that there may be alternative forms of within-host interaction (e.g., one parasite altering the infectiousness of another [5, 31]), but here we just focus on interactions affecting host susceptibility.

Host births were locally density-dependent, following a logistic model based on the total number of individuals in the patch (). Individuals died with probability *μ*_*i*_ for host class i, and we assumed infection by either parasite increased host mortality compared to uninfected hosts (i.e., *μ*_*F*_, *μ*_*B*_ and *μ*_*c*_ > *μ*_*S*_). For simplicity we primarily assumed that mortality of co-infected hosts was determined by the most virulent parasite with which they were infected [33], and so was the maximum of the mortality rates of the two single-infected hosts (i.e., *μ*_*C*_ = *max*(*μ*_*F*_, *μ*_*B*_)). We did though explore a scenario assuming additive mortality of co-infected hosts (i.e., *μ*_*C*_ = *μ*_*F*_ + *μ*_*B*_), but found this made little difference to the results (Figure S2).

Hosts were assumed to recover from infection at rate *γ*_*i*_ from infected class *i*. Recovery from single infections returned hosts to the completely susceptible state. Recovery from co-infection resulted in transition to one of the single-infected classes (either *I*_*F*_ or *I*_*B*_, depending probabilistically on the relative magnitudes of *γ*_*CF*_ and *γ*_*CB*_, denoting the recovery rates of co-infecteds from the focal and background parasites respectively).

Double-recoveries, from co-infected back to the fully susceptible class, did not happen in a single timestep. All transitions resulting in losses of individuals per host class per time step, due to infection, recovery and mortality, were modelled as a multinomial process (see Supplementary Information for details).

At the end of each timestep, the numbers of individuals of each class were subject to a movement phase. In our linear network of patches (Figure 2b), individuals could only disperse to immediate neighbouring patches each timestep. The number of individuals of class *i* moving to a neighbouring patch was determined by a multinomial draw, given the current number of individuals of that class within the source patch and the class-specific *per capita* movement probability *p*_*move,i*_. We allowed infected hosts to have reduced movement probabilities compared to uninfected hosts (i.e., *p*_*move,F*_ and *p*_*move,B*_ < *p*_*move,S*_), to reflect energetic losses or inhibition of movement behaviour due to infection. For simplicity, we assumed the movement of co-infected hosts was determined by the most debilitating parasite they were infected with (i.e., *p*_*move,C*_ = *min*(*p*_*move,F*_, *p*_*move,B*_)).

### Co-infection and co-circulation interactions

We explored three types of interaction arising from the presence of the background parasite on the focal parasite: (1) **within-host:** prior infection with the background parasite altered host susceptibility to subsequent infection by the focal parasite; (2) **demographic:** infection with the background parasite increased host mortality rate, affecting host demography and abundance of potential hosts for infection by the focal parasite; (3) **host movement:** infection with the background parasite lowered host movement rates between patches. For the within-host interaction, we varied the transmission rate of the focal parasite on background-infected hosts (*β*_*BF*_) relative to the baseline transmission rate of the focal parasite on completely susceptible hosts (*β*_*F*_). Specifically, we defined 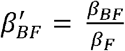, such that 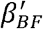 represents the relative effect of the background parasite on host susceptibility to the focal parasite; 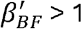 implies facilitation (hosts infected by the background parasite were more susceptible to the focal parasite than completely susceptible hosts were), whereas 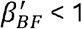 implies antagonism (hosts infected by the background parasite were less susceptible to the focal parasite than completely susceptible hosts were).

For the demographic interaction, we let the background parasite alter mortality of infected (and co-infected) hosts, by varying 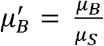, such that 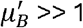 implies the background parasite was highly virulent. Note that, since the mortality of co-infected hosts was equal to the maximum virulence of either single-infected host, 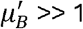 also results in a high loss rate of focal-infected hosts via increased co-infection mortality (μ_*C*_ ≫ 1).

Finally, for host movement, we let the background parasite alter between-patch movement of background-infected and co-infected hosts by varying 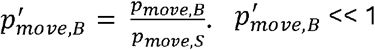 implies the background parasite imposed a high energetic constraint on infected and co-infected hosts, greatly reducing their movement between patches. Again, since we assumed the movement of co-infected hosts was equal to the minimum of either single-infected host, 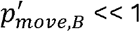 potentially restricted the spread of the focal parasite across the network. For a summary of the proximate consequences of the three interaction mechanisms for the focal and background parasites see Table S1.

### Simulation set-up and analyses

Our primary interest was to explore how these different interaction scenarios affected the spatial spread of an introduced focal parasite. All analyses involved a linear network of 20 patches. We seeded all patches with susceptible individuals at patch-level carrying capacity *K*= 1000, and introduced 20 individuals infected with the background parasite into the first patch of the chain. We then assumed two alternative ‘epidemiological scenarios’ for introduction of the focal parasite: (1) simultaneous introduction with the background parasite (i.e., co-invasion of both parasites into a fully naïve host population); or (2) introduction after the background parasite had reached a steady-state distribution across the network (timestep 200), mimicking invasion of the focal parasite into a host population infected by an endemic background parasite.

For both scenarios, we explored each of the three ways in which the background parasite may interact with the focal, as described above. Specifically, we varied either 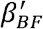 from 0 to 2 (within-host interaction), 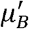 from 2 to 1 0 (demographic interaction) or (3) 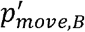 from 1 down to 0.1 (host movement). For simplicity, we explored each of these interaction effects in isolation. Other than these potential interactions, we assumed that both parasites were epidemiologically identical (i.e., both parasites used the same baseline parameter values; Table 1).

We ran the model 50 times for each parameter value and epidemiological scenario, for 1000 timesteps (initial simulations showed the system generally reached steady state by timestep 800, except under some extreme interaction scenarios; Figure S1). In what follows we primarily focus on equilibrium results (i.e., over a sampling period from timesteps 800-1000; Figure S1 regions marked (i)), although we also report results for a more narrow sampling period earlier in the simulation (timesteps 100-1 20 after introduction of the focal parasite; Figure S1 regions marked (ii)) to capture transient dynamics. In each case we report three epidemiological metrics capturing the spatiotemporal dynamics of the focal parasite: (1) the mean overall prevalence of the focal parasite (*I*_*F*_ + *I*_*C*_) across the network (*i*.*e*., the sum of all *I*_*F*_ + *I*_*C*_ hosts across all patches in the network, expressed as a proportion of the total number of hosts, averaged over the sampling period); (2) the mean across-patch coefficient of variation (CV) in prevalence of the focal parasite (*i*.*e*., the standard deviation/mean prevalence of *I*_*F*_ + *I*_*C*_ hosts across patches, averaged across the sampling period), as a measure of the degree of between-patch heterogeneity in prevalence of the focal parasite across the network; (3) the time taken for the focal parasite to establish (be present for at least 1 0 consecutive timesteps) in the final patch of the network, as a measure of its rate of spatial spread.

### Sensitivity analysis

We performed statistical analyses on the simulation data, assessing the effects of a given interaction parameter and a given epidemiological scenario (simultaneous vs delayed arrival of the focal parasite) on focal parasite success. Thus we performed analyses of covariance (ANCOVA), with overall focal prevalence, CV of focal prevalence, or last-patch focal arrival time as response variable; in a factorial model, epidemiological scenario was taken as explanatory factor and interaction parameter as a covariate (always with a linear and second-order component to account for non-linear effects). The relative importance of a given term in the statistical model was quantified as the proportion of variance explained by this term (SS_term_ / SS_total_). Data for the three metrics were z-score standardised prior to analysis. In this way, we carried out 9 ANCOVAs (3 epidemiological metrics x 3 interaction parameters).

## Experimental methods

### Biology ofthe system and interspecific interaction between parasites

*Holospora undulata* and *H. obtusa* are obligate bacterial parasites of the protist Paramecium caudatum, with similar infection life cycles but infecting different host compartments, the micronucleus and macronucleus, respectively [34]. Horizontal transmission occurs when hosts divide or die, releasing infective stages that are picked up by other *Paramecia* while feeding. In the nucleus, infectious forms differentiate into reproductive forms, capable of within-nucleus multiplication and vertical transmission in mitotically dividing hosts [35]. *Holospora* species are found worldwide in temperate locations [36], but little is known about ecological or evolutionary dynamics in natural populations [34, 37]. Local levels of infections are typically very low, even though occasional outbreaks have been reported, sometimes involving the successive arrival and replacement of species (H.D. Görtz, pers. comm.).

In the laboratory, host density can be tracked by counting the number of *Paramecium* in small samples under a dissecting microscope; infection prevalence can be tracked by fixing >20 individuals [38] and checking infection status at 1000x magnification under phase contrast. When co-circulating in experimental populations, the two parasites rarely co-infect the same host individuals (Figure S4) and tend to competitively exclude each other at the single-population level [32]. Primary exposure to H. undulata affects subsequent infection probability with the same parasite species [39]. We find here a similar effect between parasite species in a co-inoculation experiment (see Supplementary Information 2.2 for details), showing that infection success of *H. undulata* was diminished by the presence of transmission stages of *H. obtusa* (Figure S5). This suggests a within-host interaction equivalent to *β*′_*BF*_ < 1 in our model.

### Landscape experiment

Analogous to the simultaneous-release scenario in our model, we investigated the epidemic spread of the two parasites, alone and in co-circulation, for linear-chain landscapes of 5 interconnected microcosms, linked by rubber tubing (length: 5 cm, diameter: 0.8 cm) with clamps to prevent dispersal between timed dispersal events (Figure 4a). *Paramecium* were introduced into Patch 1, from where infected and uninfected hosts dispersed into the landscape. The parasite is immobile and travels together with infected hosts.

In what follows we consider *H. undulata* as the focal parasite and *H. obtusa* as the background parasite (in the Supplementary Information we present analogous results assuming roles are reversed). We carried out two experimental treatments: (i) single-parasite treatment, where 1500 uninfected hosts plus 100 hosts infected by focal parasite were introduced in Patch 1, in a total volume of 30 mL; (ii) mixed-parasite treatment, where we simultaneously introduced 1500 uninfected hosts plus 100 focal-parasite infected hosts and 100 background-parasite infected hosts into Patch 1. Every treatment had six landscape replicates, organised in two experimental blocks (shifted in time by two days). Results for the corresponding single background-parasite treatment are shown in the Figures S8-S9, and for an additional control treatment (1600 uninfected hosts in Patch 1) in Figure S8.

By removing the connection-blocking clamps, free dispersal between patches was allowed for three hours, three times a week, for a total of four weeks. Under these conditions, uninfected hosts typically reach new patches one to several time steps ahead of the parasite (Figure S6; [40]), such that dynamics closely resemble spreading epidemic waves. Prior to every second opening of connections, we quantified *Paramecium* density and infection prevalence in each patch.

### Statistical analysis

We analysed spatial epidemic spread rate as a “front patch progression” [40] by way of linear regression, with parasite front patch position (*i*.*e*., farthest downstream patch with infection detected, ranging from 1 to 5) as response variable and time (counted as the number of between-patch opening events) as a covariate. Parasite spread rate corresponds to the slope of this regression. In a factorial model, we included parasite treatment (single vs mixed) and added landscape identity and experimental block as random factors (LMM).

Final infection prevalence (proportion of infected individuals in a patch) was analysed using a Generalised Linear Mixed Model (GLMM) with binomial error. Parasite treatment and patch position were included as explanatory factors, and landscape identity as a random factor. We further calculated the among-patch coefficient of variance (CV) in final infection prevalence (mean / standard deviation) for each landscape, which was then compared between parasite treatments.

## RESULTS

### Epidemiological model results

#### 1) Within-host interaction (*β*′_*BF*_)

Overall, the co-infection interaction affecting host susceptibility (*β*′_*BF*_) had the greatest impact out of all interaction scenarios on each of the three epidemiological metrics of the focal parasite (Figure 3). Analysis of variation in simulation outcomes of the equilibrium dynamics shows that variation in *β*′_*BF*_ explained a large part of the observed variation in all three epidemiological metrics (63-99%; Table S2). When prior infection by the background parasite completely inhibited subsequent infection by the focal (*β*′_*BF*_ = 0), the mean network prevalence of focal-infected hosts was ∼2% when introduced at the same time as the background parasite. At the highest interaction value examined (*β*′_*BF*_ = 2, such that background-infected hosts were twice as likely as uninfected hosts to be infected by the focal parasite), the mean focal-infected prevalence was ∼65% (Figure 3b).

**Figure 3.**
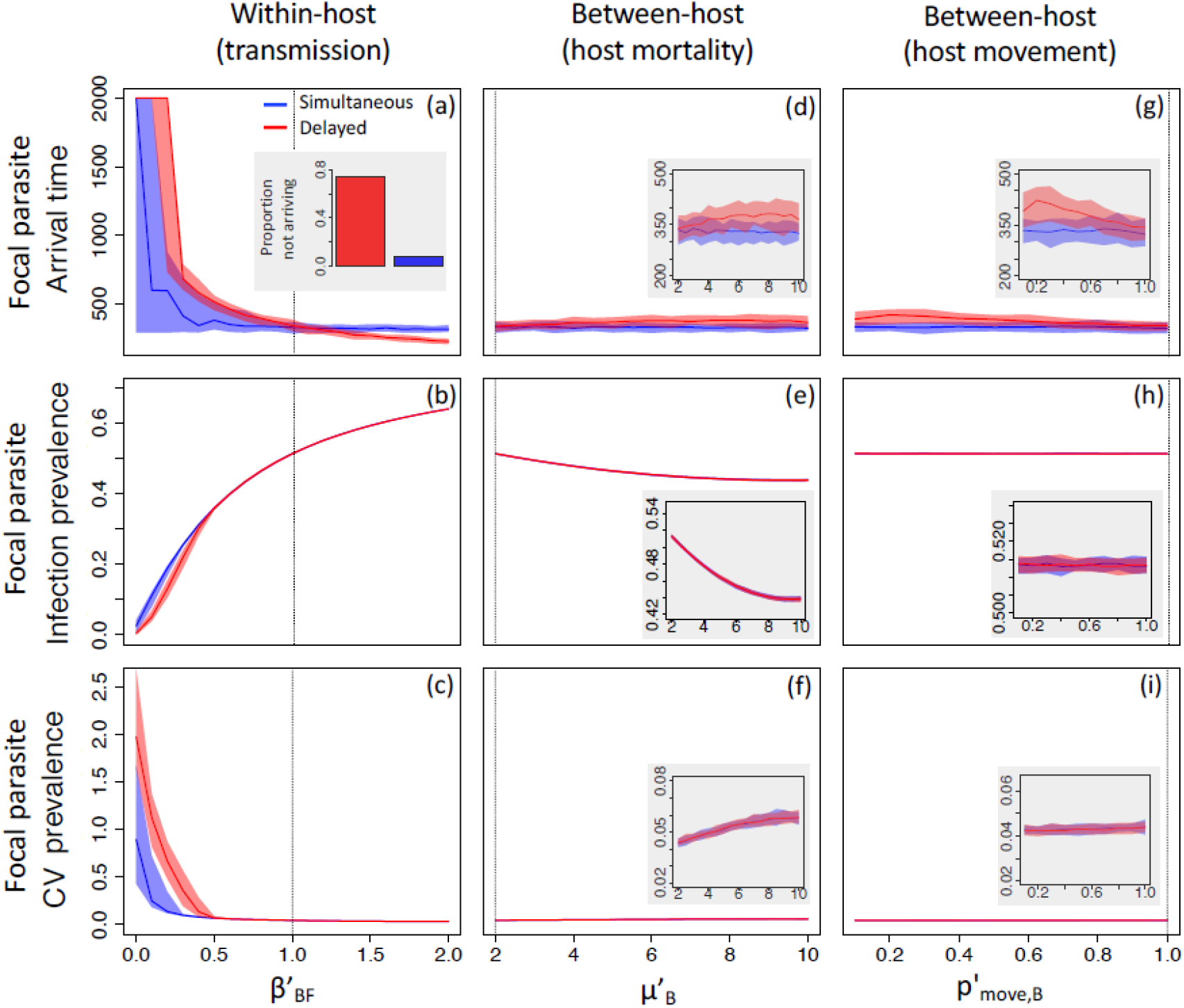
Model output, showing the effects of three types of parasite interactions by the background parasite on the spatial spread of the focal parasite at equilibrium, under two co-circulation scenarios (simultaneous introduction or delayed introduction of the focal). **(a)-(c)**: background parasite alters host susceptibility to the focal parasite via changes in *β*′_*BF*_; **(d)-(f)**: background parasite increases host mortality via changes in *μ*′_*B*_; **(g)-(i)**: background parasite reduces host movement probability via changes in *p*′_*move,B*_). Epidemiological metrics for the focal parasite relate to the time to spread from Patch 1 to the end patch of the network (top row), mean landscape-wide infection prevalence (middle row) and mean between-patch coefficient of variation of prevalence (standard deviation/mean; bottom row). Solid lines show the median values of 50 replicated simulations for each parameter value, shaded areas show the 95% quantiles of those values. The vertical dotted lines in each panel show the baseline (non-interaction) parameter values in each case. Small insert graphs in panels d-i show zoomed-in versions of the results. The insert graph in panel a shows the proportion of replicate runs with the focal parasite failing to reach the end patch, following simultaneous or delayed introduction (calculated across simulations for 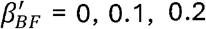; the focal parasite reached the end patch in all simulations where 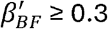).

Importantly, the impact of prior infection by the background parasite on host susceptibility to the focal affected the spatial dynamics of the focal parasite across the network. Most notably, at lower values of *β*′_*BF*_, the focal parasite did not always reach the end point of the network, and this effect was amplified if the focal parasite was introduced after the background parasite had established (Figure 3a inset figure). At the extreme, *β*′_*BF*_ = 0, the probability of the focal parasite reaching the end-patch of the network was 20% when introduced at the same time as the background parasite, whereas it did not reach the final patch in any of the simulations where it was introduced after the background parasite had first established itself across the whole network. Hence under very strong interactions affecting host susceptibility, the model predicted a patch-level priority effect, whereby occupation of a patch by the background parasite limited the ability of the focal parasite to establish in that patch. This was particularly apparent if the background parasite was allowed to establish first, limiting the ability of the focal parasite to invade and spread across the network.

Furthermore, even if the focal parasite was able to persist, strong negative interactions via host susceptibility (*i*.*e*., reducing *β*′_*BF*_) increased the time taken for the focal parasite to establish in the final patch of the network (Figure 3a) and spatial heterogeneity of focal parasite prevalence across the network (as measured by the between-patch coefficient of variation (CV) in focal parasite prevalence; Figure 3c). These effects were generally magnified if the background parasite was allowed to establish before the focal parasite invaded; analysis of variance shows ∼1 0% of the variation in CV and end-patch arrival time were explained by an interaction between p’_*BF*_ and epidemiological scenario (Table S2). Hence a co-infection interaction, whereby the background parasite impacts individual-level susceptibility to the focal, results in changes both to overall prevalence of the focal parasite across the network, and also to the probability and rate of spatial spread of the focal, and to its spatial distribution (heterogeneity) across the network. In particular, if there is a strong co-infection interaction (e.g., *β’*_*BF*_ = 0.2; Figure S1 a) there is very high heterogeneity in focal prevalence across the network, as patches further down the chain do not reach their equilibrium values within the sampling window (the grey zone marked (i) in Figure S1). Hence, strong co-infection interactions limit dispersal, slowing establishment rate in remote patches, leading to high levels of heterogeneity across the network. These spatial effects can be magnified if the background parasite is able to establish in the landscape before the focal parasite invades. For facilitative interactions (*β*′_*BF*_ > 1), we observed little effect on our summary statistics, likely due to dispersal becoming the rate limiting factor. In particular, there was little additional effect past *β*′_*BF*_ > 1 on accelerating arrival of the focal parasite in the end-patch of the network (Figure 3a) or reducing between-patch CV in focal prevalence (Figure 3c).

#### 2) Between-host interactions (μ’_B_ and p’_move,B_)

In general, effects due to the other interaction parameters (via host mortality, *μ’*_*B*_; Figure 3d-f, and host movement, *p*′_*move,B*_; Figure 3g-i) were much weaker in comparison to the within-host interaction (*β*′_*BF*_). Analysis of variation suggested that host mortality due to prior infection by the background parasite, *μ*′_*B*_, explains most of the variation in mean and CV of focal infection prevalence (>85%; Table S2). Increasing *μ*′_*B*_ resulted in a reduction in prevalence of the focal parasite (Figure 3e), albeit with little apparent effect on the CV in focal prevalence across the network (Figure 3f; although close inspection suggests there was a marginal trend for increasing *μ’*_*B*_ to marginally increase CV of the focal parasite; Figure 3f inset).

Interactions involving the probability of host movement (*p*′_*move,B*_) had no discernible effect on overall mean prevalence (Figure 3h) or CV of prevalence (Figure 3i) of the focal parasite across the network (analysis of variation suggests variation in *p*′_*move,B*_ explains <8% of the variation in these metrics; Table S2).

There were, though, marginal effects of both *μ*′_*B*_ and *p*′_*move,B*_ on the time taken for the focal parasite to spread to the end-patch of the network, but largely only when it invaded after the background parasite had established (Figure 3d, Figure 3g red lines). Analysis of variation suggested variation in *μ’*_*B*_ and *p*′_*move,B*_ explained ∼8% and ∼20% of the variation in patch arrival (main effects + interactions) respectively, although these effects were both influenced by interactions with epidemiological scenario, which alone dominated the amount of variation in end-patch arrival time (main effects of epidemiological scenario of 23% and 35% respectively; Table S2). For both interaction parameters, the stronger the interaction (i.e., the higher *μ*′_*B*_ or the lower *p*′_*move,B*_), the longer it took for the focal parasite to spread and establish in the end-patch of the network. Again, these effects were not seen when the focal parasite was introduced at the same time as the background parasite (Figure 3d, Figure 3g blue lines). Hence, demographic or movement-mediated interactions can slow progression of an invading pathogen into a system with an established background parasite, but not when the two invade concurrently. Together, these analyses show that each interaction type leaves a characteristic signature across the different epidemiological metrics, with the within-host interaction (*β*′_*BF*_) causing the most consistent impact overall.

We note that the above results relate to the equilibrium phase of the dynamics (i.e., for timesteps 800-1000 of simulations). Qualitatively similar results were found for the transient phase (over timesteps 100-1 20 after introduction of the focal parasite), although overall prevalences were lower and the degree of between-patch heterogeneity was higher during the transient phase (Figure S3). Furthermore, the focal parasite did not reach the end-patch of the landscape during the transient phase.

### Experimental results

Once introduced in Patch 1 of the landscapes, the focal parasite (H. undulata) produced a series of local outbreaks in downstream patches, with infection levels reaching up to 85% (Figures S7, S8). Simultaneous addition of the background parasite (*H. obtusa*) slowed the spatial spread of the focal parasite (Treatment *x* time: *X*^2^_1_p = 0.008; Table S3). This is visualised in Figure 4b, where the progression of the = 6.98, epidemic front in the co-circulation treatment has a shallower slope over time than in the single-parasite treatment. Thus, while the focal parasite arrived in the last two patches of the landscape (4 and 5) in five out of six replicates of the single-parasite treatment, it failed to get past patches 2 or 3 in three out of six replicates of the co-circulation treatment.

**Figure 4.**
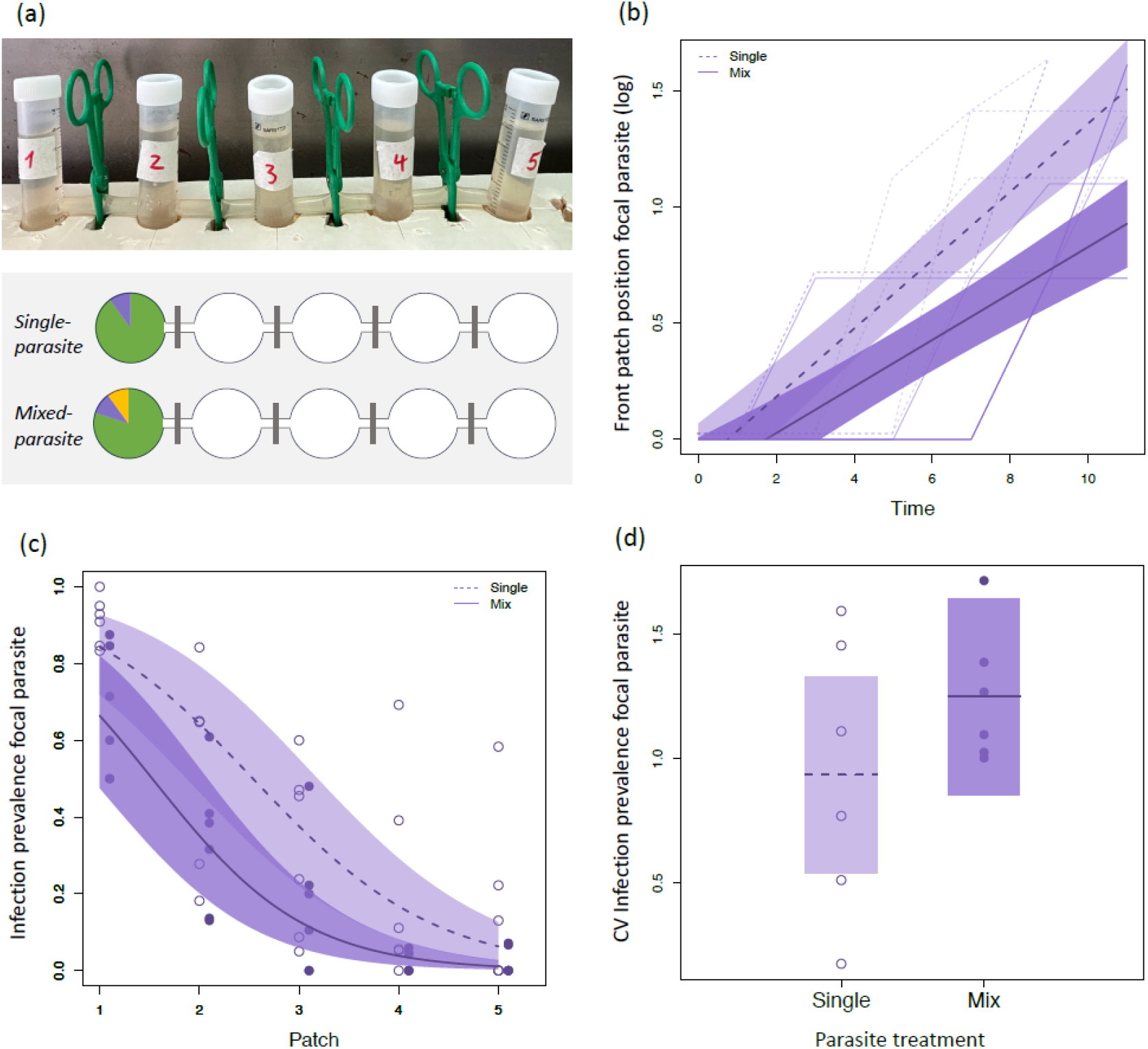
Experimental results. **(a)** Design of co-circulation experiment, with single-parasite (focal parasite, *Holospora undulata*) and mixed-parasite treatments (focal parasite and background parasite, *H. obtusa*). Experimental landscapes were composed of 5 tubes interconnected by silicon tubing (photo). Connection-blocking clamps were removed 3x a week for 3h allowing the natural dispersal of hosts. **(b)** Rate of spatial spread of the focal parasite (*H. undulata*) into the 5-patch landscapes, in single-parasite and mixed-parasite treatments. The front position is plotted against time (= number of 3h opening events), and thus the slope of the relationship describes the rate of spread. **(c)** Infection prevalence of the focal parasite across the 5 patches in the landscapes, in single-parasite and mixed-parasite treatments. **(d)** Coefficient of variation in final prevalence of the focal parasite (mean/SD across the 5 patches in a landscape). The graphs show raw data (thin lines and points) as well as model predictions for the two treatments (mean and 95% range).

The presence of the background parasite also led to significantly lower final average infection prevalences of the focal parasite (*X*^2^ _1_= 7.28, p = 0.006; Table 3; Figure 4c). On average, its landscape-wide prevalence was reduced by nearly half in the co-circulation treatment (0.24 ± 0.03 SE), compared to the single-parasite treatment (0.43 ± 0.05). This difference still holds when omitting cases of non-colonisation and considering only the first two patches, occupied by both parasites in all landscapes (*X*^2^ = 7.57, p = 0.021).

However, spatial variation in final prevalence (among patches of the same landscape) did not significantly differ between treatments (CV prevalence F_1,1 0_ = 1 .56, p > 0.2; Figure 4d).

## Discussion

While it is well known that parasite-parasite interactions can alter host-level infection processes (host susceptibility, disease progression etc [7,8,10]), with growing recognition of potential impacts of those interactions for host and parasite population dynamics [3,28,41,42], we have little understanding of their consequences for parasite epidemiology across landscapes. Combining stochastic simulation models with lab infection experiments, we show that different forms of parasite-parasite interactions can impact parasite spatiotemporal dynamics in different ways. These potentially result in patch-level priority effects that alter parasite spread across the landscape, particularly if parasites are introduced sequentially to the host population.

### Simulation model results: within-host interaction has the strongest effect on co-circulation dynamics

Our model showed that all three interaction types explored (within-host interaction affecting host susceptibility; demographic interaction affecting host survival; movement interaction affecting host dispersal probabilities) altered the spatiotemporal dynamics of the focal parasite (the species being impacted by the other, background parasite). Dominant among these effects was the impact of the within-host interaction on host susceptibility (via changes in the transmission parameter 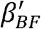). Varying this parameter had strong effects on the rate of spread of the focal parasite (Figure 3a), its landscape-level prevalence (Figure 3b) and its spatial heterogeneity across the landscape (Figure 3c). Importantly, these spatial effects were considerably amplified when the focal parasite invaded after the background parasite had established across the landscape.

In contrast, we found that effects due to the other two forms of interaction (demographic and movement related) were considerably weaker than the within-host interaction, with the movement interaction in particular having little detectable effect on landscape-wide prevalence (Figure 3h) or spatial heterogeneity (Figure 3i) of the focal parasite. Both these interactions, though, did impact the rate of spread of the focal parasite across the landscape (Figures 3d,g). Notably, these impacts were only apparent when the focal parasite invaded after the background parasite had established across the landscape. Contrastingly, effects on rate of spread due to the within-host interaction were generally seen regardless of whether the focal parasite was introduced after or at the same time as the background parasite (Figure 3a).

Arguably, this dependence (or not) of an effect on the relative timing of arrival could help distinguish between interactions that are mediated via changes in host density or reduced infectivity, although the magnitudes of those effects were marginal in our simulations. Regardless, the epidemiological consequences of parasite-parasite interactions can be amplified by priority effects acting across the landscape, when the dominant parasite species is allowed to establish in a patch prior to invasion by the impacted species.

The dominance of within-host interactive effects in our analysis likely stems from the direct impact of the background parasite on transmission success of the focal, and the lack of cost for the background parasite itself. For the two between-host interaction types (increased mortality, reduced dispersal), the background parasite is also affected detrimentally, whereas the focal parasite is only directly affected in co-infected hosts (summarised in Table S1). Indeed, under a dispersal interaction, the focal parasite may even have an advantage as background-infected hosts disperse less overall than the focal parasite, potentially allowing the focal to establish in neighbouring patches, in single-infected hosts, ahead of the background parasite. There may be some support for this from the experimental system, where we see a generally lower spread rate of the background parasite. This may be related to a differential effect on host dispersal, but also simply reflect the lower local prevalences (and hence lower dispersal probability). Furthermore, the between-host interactions will alter local demography, which may be compensated for by within-patch density-dependent processes. For example, high mortality of co-infected hosts will remove focal-infected hosts from the population, but will also allow more susceptible hosts to be born due to density-dependent release (e.g., [5]), reducing the net impact on the focal parasite.

### Co-circulation in experimental landscapes: landscape-level priority effects and the role of stochasticity

We sought support for some of our predictions through a series of experiments with our biological system. In that system, the background parasite (*H. obtusa*) reduces infection success of the focal parasite (*H. undulata*; Figure S5), similar to the within-host interaction mechanism in the model (i.e., 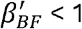). We tested the performance of the focal parasite across a spatial landscape of connected patches under single-parasite and co-circulation treatment experiments, with simultaneous introduction of both parasites in the start patch. As predicted by the model (Figure 3a-c), the presence of the background parasite led to a lower rate of spread of the focal parasite, such that it frequently failed to reach the most distant patches (Figure 4b). We also, as predicted, found a lower landscape-wide infection prevalence of the focal parasite (Figure 4c).

Our full landscape experiment did not specifically test model predictions relating to the importance of sequential parasite invasion. However, preliminary tests exploring sequential scenarios under simplified spatial settings are consistent with outcomes in our model (see Supplementary Information 2.4 for details). In one experiment, we transferred infected hosts into a single uninfected population with varying delays between introduction of the background and focal parasites. We found strongly dampened outbreaks of the focal parasite when it was introduced one week after the background parasite and, when introduced with a 2-week delay, the focal parasite failed to establish (Figure S1 1). Similar results were obtained in 2-patch interconnected tubes, when the focal parasite was allowed to disperse naturally from one patch into a second patch already infected with the background parasite (Figure S1 2). These observations are consistent with the general idea that multiparasite dynamics can play out on a first-come, first-served basis (i. e., priority effects [9, 42-46]), but acting at the patch- or even metapopulation-scale. Similar effects also exist in medically relevant diseases [47], for example where cross-immunity can create local priority effects, protecting endemic (resident) parasite variants or species against invasion by newly arriving parasites [27].

In our model, within-host interactions with the background parasite occur across the entire landscape, thus limiting spatial spread of the focal parasite. Yet, in our experimental landscapes, the background parasite itself showed limited spatial expansion, in part due to competitive effects from the focal parasite (Figure S9). This raises the question of how the focal parasite’s spread was affected in downstream patches if the background parasite was not detected in those patches. We hypothesise that upstream effects propagated into the landscape in two possible ways. First, there is evidence that (cross-)resistance can be induced upon contact with these parasites ([39]; J. Zabalegui unpubl. data), without causing infection. Thus, contact with the background parasite may increase resistance in upstream patches, and primed uninfected hosts could then colonise the more distant patches. Consequently, the focal parasite will be confronted with more resistant hosts and experience reduced spread rates. This hypothesis could be tested by comparing the resistance of downstream hosts from the different treatments (control, single-, mixed-parasite).

Another possibility relates to stochastic parasite diffusion dynamics. Due to small infected population sizes and low host dispersal, “parasite jumps” to a new downstream patch were a rare event in our experiments (mean 0.44 *H. undulata* jumps per opening of connections, across all treatments). This stochastic effect may be compounded by competition with the background parasite, further reducing the number of hosts infected with the focal parasite, and, consequently, reducing patch colonisation and spatial spread rate. Such a delayed spread read could be initiated in the upstream patch(es) and then cascade into the landscape. Consistent with this idea, we found a significant positive relationship between focal-infected density in one patch at a given time point, and the probability of detection of infection in the next patch in the chain at the next time point (*X*^2^ _1_ = 5.1, df = 1, p = 0.024; GLM of density log-transformed and patch position as a factor, all data combined; see also [40]). These observations match previous theory that, at the individual patch level, stochastic effects acting on small population sizes can restrict pathogen establishment and growth, even if R_0_ > 1 [49]. At the metapopulation level, these effects can amplify, such that low numbers of dispersers can restrict parasite establishment in successive patches, limiting overall parasite spread and persistence [50]. To allow for stochastic and/or transient effects in our simulations, we implemented a low dispersal probability (1% of hosts dispersed between patches on average per time step). Accordingly, we see high variation in arrival time when within-host competition is strong, but this rapidly diminishes as the background parasite becomes facilitative and dynamics become more deterministic (Figure 3c). In contrast, altering movement probability had little effect on outcome variation.

In the context of co-circulating parasites, the impact of these stochastic spatial effects might be additionally modulated by the relative abundances of the two parasite species. This is seen in our experiments, where we varied the relative numbers of focal and background hosts in a single patch (Figure S5) or between patches (Figures S11, S1 2). Successful focal parasite invasion was more likely when either arriving in large numbers or when resident background infection levels were low. At the wider landscape level, the epidemiological consequences of parasite co-circulation will therefore depend on the interaction between local abundances, host dispersal rates and stochastic forces.

### Interactions between interactions

For simplicity and clarity, our model assumed both parasite species were epidemiologically identical (same baseline transmission, virulence, recovery and dispersal rates), apart from a given interaction effect tested. In reality, parasite species or strains may differ in baseline epidemiological parameters, which could alter spatiotemporal outcomes. In our experimental system, for example, focal and background parasites differ in their effects on host population growth and possibly also on dispersal (J. Zabalegui, unpubl. data). Poletto et al [18] showed theoretically that if co-circulating parasites differ in infectious period, this can interact with dispersal rates to alter spatiotemporal outcomes. Furthermore, we considered each interaction type in isolation, whereas multiple pathways of interaction between parasites, including bi-directional (focal affecting background and vice versa), could occur. For example, while our experiments showed strong effects of H. obtusa (our background parasite) on H. undulata (our focal), we did find evidence of effects, albeit weaker, in the opposite direction (H. undulata reducing prevalence of H. obtusa; Supplementary Information 2.3.3). Incorporating these multiple interaction pathways, could greatly increase the complexity of potential outcomes, so it would be valuable to extend our analyses to capture the likely richness of interaction scenarios occurring in the real world.

### Conclusions

Overall, we show that parasite interactions can scale up to affect the spatiotemporal epidemiology of co-circulating parasite species, but those effects can differ, depending on the type of interaction occurring and the epidemiological scenario (i. e., the timing of invasion of the parasite species relative to each other into the host population). Both our simulation and experimental results suggest that within-host parasite interactions which alter host susceptibility to infection can have particularly strong effects on the rate of spatial spread and distribution of parasites across the landscape. Translating these predictions and findings to real-world disease scenarios would require more in-depth analyses of potentially multiple interaction and epidemiological scenarios.

Nevertheless, given the ubiquity of co-circulating parasite species in natural host populations, and the known occurrence of strong interspecific interactions between them, our findings highlight the importance of understanding these overlooked processes to better inform forecast models and disease management strategies in the future.

## Supporting information

Simulation details and supplementary experiments

## ACKNOWLEDGEMENTS

AF and ADD would like to thank the Natural Environment Research Council (NERC; grant ref: NE/X01424X/1). OK was supported by a grant from the French Agence Nationale de la Recherche (ANR-20-CE02-0023-01). This is publication ISEM-202X-XXX of the Institut des Sciences de l’Evolution, Montpellier.

