## Supplementary material for "Differing effects of parasite-parasite interaction types on the spatial epidemiology of co-circulating parasites": Simulation details and supplementary experiments

### 1) Additional simulation model details

#### *1.1 Within-patch dynamics*

Our model follows a stochastic Monte Carlo formulation, tracking changes in the integer numbers of individuals of each the following classes within each patch: $S$ (susceptible hosts, not currently infected by either parasite), $I_F$ (infected by the focal parasite only), $I_B$ (infected by the background parasite only), and $I_C$ (simultaneously co-infected by both parasites). The model iterates on a fixed time step (Δt = 1), processing the dynamics occurring within each patch each time step in the following order: host births; infection gains and losses and host mortality; movement.

Host reproduction was assumed to be locally density dependent, following a logistic model based on the total number of individuals in the patch ($N_i = S_i + I_{F,i} + I_{B,i} + I_{C,i}$, where *i* denotes the current patch), such that the number of births in the current patch at the current timestep followed a Poisson process with mean $bN_i\left(1-\frac{N_i}{K'}\right)$, where *b* is the per capita host birth rate and $K'$ determines the strength of density dependence acting on host reproduction (such that the per patch carrying capacity in the absence of disease, $K = K'\left(\frac{b-\mu_S}{b}\right)$, where $\mu_S$ is the susceptible host mortality rate). Note, all host classes were assumed to contribute equally to reproduction (no impact of infection status on reproduction rate), and all births were assumed to be uninfected by either parasite (no vertical transmission).

The infection process was assumed to follow a frequency-dependent transmission process, based on the numbers of susceptible, infected and total hosts within the current patch at the current time step (newborns were assumed not to contribute to these processes in the current time step). Specifically, the loss of individuals per host class per time step, due to infection, recovery and mortality, were modelled as a multinomial process, governed by the following transition rates (note in what follows we drop the patch-specific labelling for clarity – the following terms and definitions refer to the numbers within a given patch at a given time point):

$$S: \left\{\frac{\beta_F(I_F+I_C)}{N}, \frac{\beta_B(I_B+I_C)}{N}, \mu_S\right\}$$

$$I_B: \left\{\frac{\beta_{BF}(I_F+I_C)}{N}, \gamma_B, \mu_B\right\}$$

$$I_F:\left\{\frac{\beta_{FB}(I_B+I_C)}{N},\gamma_F,\mu_F\right\}$$

$$I_C:\{\gamma_{CB},\gamma_{CF},\mu_C\}$$

The first two transition terms from the *S* class represent loss of susceptibles due to transmission (contracting infection) from $I_F$- and $I_B$-infected hosts, respectively. Note that co-infected hosts ($I_C$) contribute to the force of infection by each parasite type on susceptible hosts, and these contributions are equal to that of single-infected hosts of that type (i.e., co-infection was assumed not to alter infectiousness of co-infected hosts relative to single-infected hosts). The $\beta_i$ terms are the baseline per capita transmission rates for parasite *i* on susceptible hosts. The $\beta_{ji}$ terms represent the transmission rate of parasite *i* on hosts already infected by parasite *j*. If these $\beta_{ji}$ terms differ from the corresponding transmission rate for that parasite on susceptible hosts (the corresponding $\beta_i$) then this allows for a within-host interaction such that prior infection by parasite *j* alters host susceptibility to parasite *i* (see below for details of these interactions). Note that we assume susceptible individuals cannot transition directly to being co-infected hosts (i.e., they cannot contract infections by both parasites) in a single timestep.

The $\mu_i$ terms represent the mortality rates of each host class *i*, and the $\gamma_i$ terms represent the recovery rates from infected class *i*. While different assumptions could be made about possible additive or multiplicative effects of co-infection on host mortality, for simplicity we assumed the co-infected mortality rate, $\mu_C$, was the maximum of the mortality rates of the two single-infected hosts (i.e., $\mu_C = max(\mu_F, \mu_B)$, such that mortality of co-infected hosts was determined by the most virulent parasite they were infected with). Recovery from single-infections return hosts to the completely susceptible state (following the SIS framework). Recovery from co-infection was assumed to result in transition to one of the single-infected classes (either $I_F$ or $I_B$, depending probabilistically on the relative magnitudes of $\gamma_{CF}$ and $\gamma_{CB}$, denoting the recovery rates of co-infecteds from parasite 1 or parasite 2 respectively); it was assumed double-recoveries (i.e., from co-infected back to the fully susceptible class) did not happen in a single timestep.

Given the above transition rates, the total number of losses from class *i* in a given timestep was determined from a binomial draw given the number of individuals of that class at the start of the timestep, and overall transition probability $1 - e^{-h_i}$ where $h_i$ was the sum of all transition rates for that class. Those events were then partitioned among the alternative transition paths according to a multinomial process, with event-specific probabilities given by the above transition rates divided by the overall rate $h_i$. Individuals transitioning out of each class due to infection or recovery then entered the appropriate destination class for the start of the next timestep (e.g., the number of susceptible individuals that were deemed to contract infection by the background parasite were removed from the *S* class and added to the $I_B$ class; the number of $I_i$ hosts recovering were added to the *S* class, etc).

#### *1.2 Between-patch movement dynamics*

At the end of each timestep, the numbers of individuals of each class remaining, after births, deaths, infections and recoveries had been accounted for, were subject to a movement phase. We assume the network is structured as a linear chain of patches (Figure 2B), such that individuals from one patch can only disperse to immediate neighbouring patches each timestep. For simplicity we assume there is no directionality to movement, such that individuals dispersing from non-terminal patches are equally likely to move to either neighbour; individuals in terminal patches (the first or last patch in the chain) can only move to the immediate neighbour (patch 2 or the penultimate patch, respectively).

The number of individuals of each class *i* moving to each available neighbouring patch was determined by a multinomial draw given the current number of individuals of that class within the source patch and the class-specific per capita movement probability $p_{move,i}$. Moving individuals were added to the relevant class lists of the corresponding neighbouring patches for the start of the next timestep. We allowed infected hosts to potentially have reduced movement probabilities compared to uninfected hosts (i.e., $p_{move,1}$ and $p_{move,2}$ < $p_{move,S}$) to reflect energetic losses or inhibition of movement behaviour due to infection. As with co-infected mortality, it is possible to envision a range of potential impacts of co-infection on host movement behaviour, but for simplicity we assumed the co-infected movement probability, $p_{move,C}$, was the minimum of the movement probabilities of the two single-infected hosts (i.e., $p_{move,C} = min(p_{move,1}, p_{move,2})$, such that movement of co-infected hosts was determined by the most debilitating parasite they were infected with).

**Table S1.** Proximate consequences of the three interaction mechanisms for the background and focal parasite species

| Interaction mechanism | Consequence for Background parasite | Consequence for Focal parasite |
|---|---|---|
| Altering host susceptibility to focal<br><br>($\beta'_{BF}$ < 1 or $\beta'_{BF}$ > 1) | None | $\beta'_{BF}$ < 1 (or $\beta'_{BF}$ > 1) reduces (or increases) number of hosts available for infection by focal |
| Increasing mortality of background-infected (single and co-infected) hosts ($\mu'_{B}$ > 1) | Reduces infected lifespan (of single and co-infected hosts), and hence onward transmission opportunities | 2-fold:<br><br>1) As for background parasite, reduces infected lifespan and onward transmission opportunities of co-infected hosts<br><br>2) Reduces number of background-infected hosts available for potential infection by focal |
| Reducing movement of background-infected (single and co-infected) hosts ($p'_{move,B}$ < 1) | Reduces potential spatial spread of background parasite (in single and co-infected hosts). At extreme ($p'_{move,B}$ → 0), may prevent spread due to stochastic losses during dispersal phase | 2-fold:<br><br>1) As for background parasite, reduces potential spatial spread via co-infected hosts<br><br>2) Potential beneficial effect by reducing spread of competitor (background parasite). Likely to only be realised if background parasite has additional adverse effect on focal (e.g., if $\beta'_{BF}$ < 1) |

**Table S2.** Sensitivity analyses (Analyses of Covariances, ANCOVA) for data from simulation models quantifying the variation in epidemiological metrics for the focal parasite, as a function of the interaction parameter and epidemiological scenario (simultaneous vs delayed introduction of the focal parasite relative to the background parasite). Separate analyses were carried out for each of three interaction types ($\beta'_{BF}$, $\mu'_B$, $p'_{move,B}$). Data for the three metrics (infection prevalence, CV of prevalence, time of focal parasite to arrive in landscape end patch) were z-score standardised prior to analysis. Factorial ANCOVA models contained the linear and quadratic terms for the interaction parameter (covariate) and introduction scenario. The proportion of variance (sums of squares, SS) is reported for each term in the model explained ($SS_{term}$ / $SS_{total}$).

| | Prevalence<br>Model $R^2$ = 0.99 | CV prevalence<br>Model $R^2$ = 0.63 | Arrival time<br>Model $R^2$ = 0.79 |
|---|---|---|---|
| $\beta'_{BF}$ | 0.87 | 0.29 | 0.51 |
| ${\beta'_{BF}}^2$ | 0.12 | 0.26 | 0.15 |
| Scenario | <0.01 | <0.01 | <0.01 |
| Scenario x $\beta'_{BF}$ | <0.01 | 0.05 | 0.10 |
| Scenario x ${\beta'_{BF}}^2$ | <0.01 | 0.04 | 0.01 |

| | Prevalence<br>Model $R^2$=0.995 | CV prevalence<br>Model $R^2$=0.86 | Arrival time<br>Model $R^2$=0.43 |
|---|---|---|---|
| $\mu'_B$ | 0.90 | 0.82 | 0.02 |
| ${\mu'_B}^2$ | 0.09 | 0.04 | 0.01 |
| Scenario | <0.01 | <0.01 | 0.23 |
| Scenario x $\mu'_B$ | <0.01 | <0.01 | 0.04 |
| Scenario x ${\mu'_B}^2$ | <0.01 | <0.01 | 0.01 |

| | Prevalence<br>Model $R^2$=0.006 | CV prevalence<br>Model $R^2$=0.09 | Arrival time<br>Model $R^2$=0.54 |
|---|---|---|---|
| $p'_{move,B}$ | <0.01 | 0.08 | 0.10 |
| ${p'_{move,B}}^2$ | <0.01 | <0.01 | <0.01 |
| Scenario | <0.01 | <0.01 | 0.35 |
| Scenario x $p'_{move,B}$ | <0.01 | <0.01 | 0.08 |
| Scenario x ${p'_{move,B}}^2$ | <0.01 | <0.01 | <0.01 |

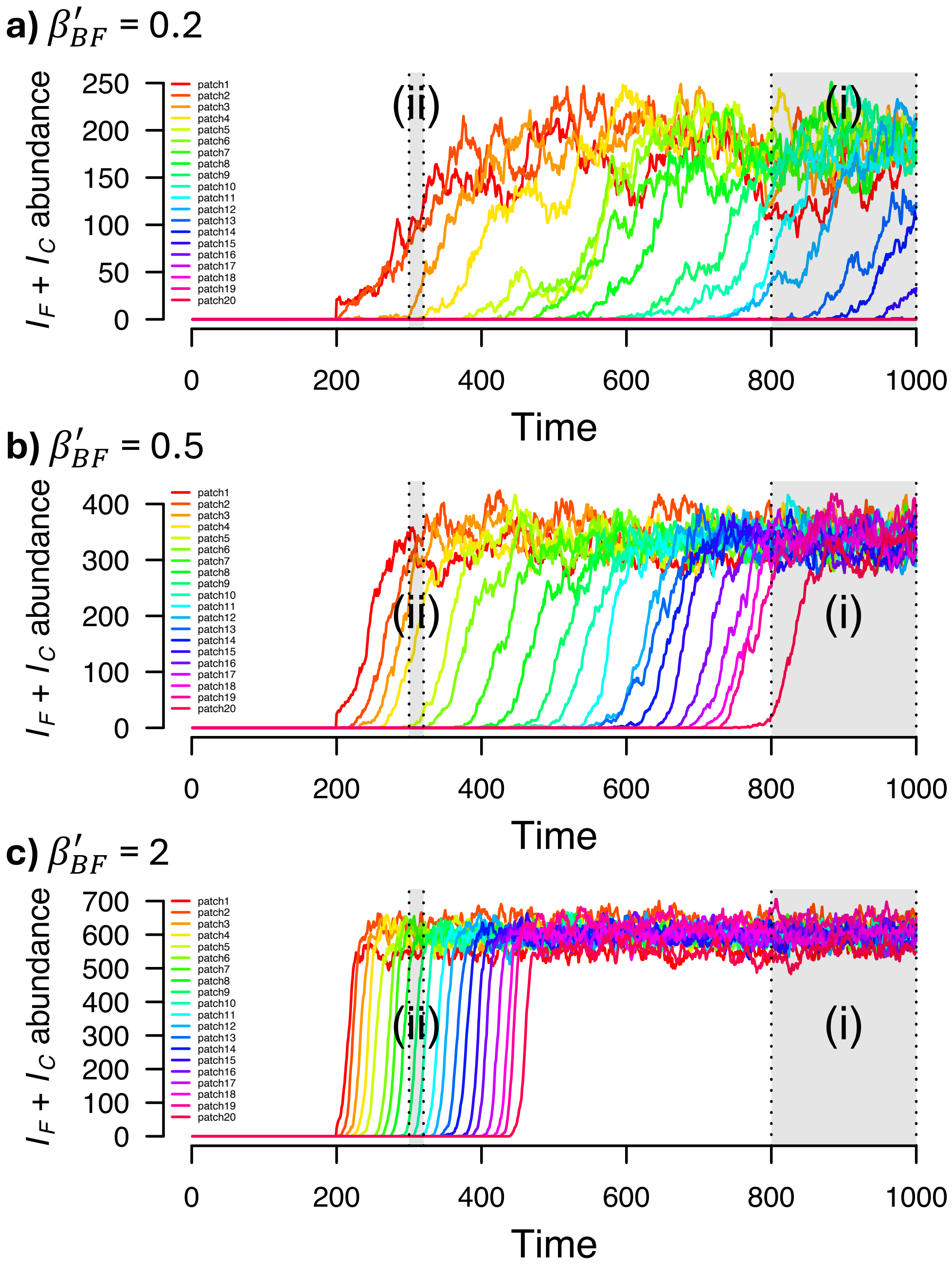

**Figure S1.** Example simulation results showing the overall abundance of the focal parasite ($I_F + I_C$) in each patch across the network over 1000 simulated time points, for different values of $\beta'_{BF}$, the effect of prior infection by the background parasite on infection probability by the focal: **(a)** $\beta'_{BF}$ = 0.2; **(b)** $\beta'_{BF}$ = 0.5; **(c)** $\beta'_{BF}$ = 2. The focal parasite was introduced into Patch 1 of the landscape at time point 200 (the background parasite was introduced into Patch 1 at time point 1). The two grey shaded regions show the sampling windows for summarising results: (i) 'equilibrium' dynamics (time points 800 to 1000); (ii) 'transient' dynamics (time points 300 to 320 (100 to 120 time steps after initial introduction of the focal parasite into the landscape)).

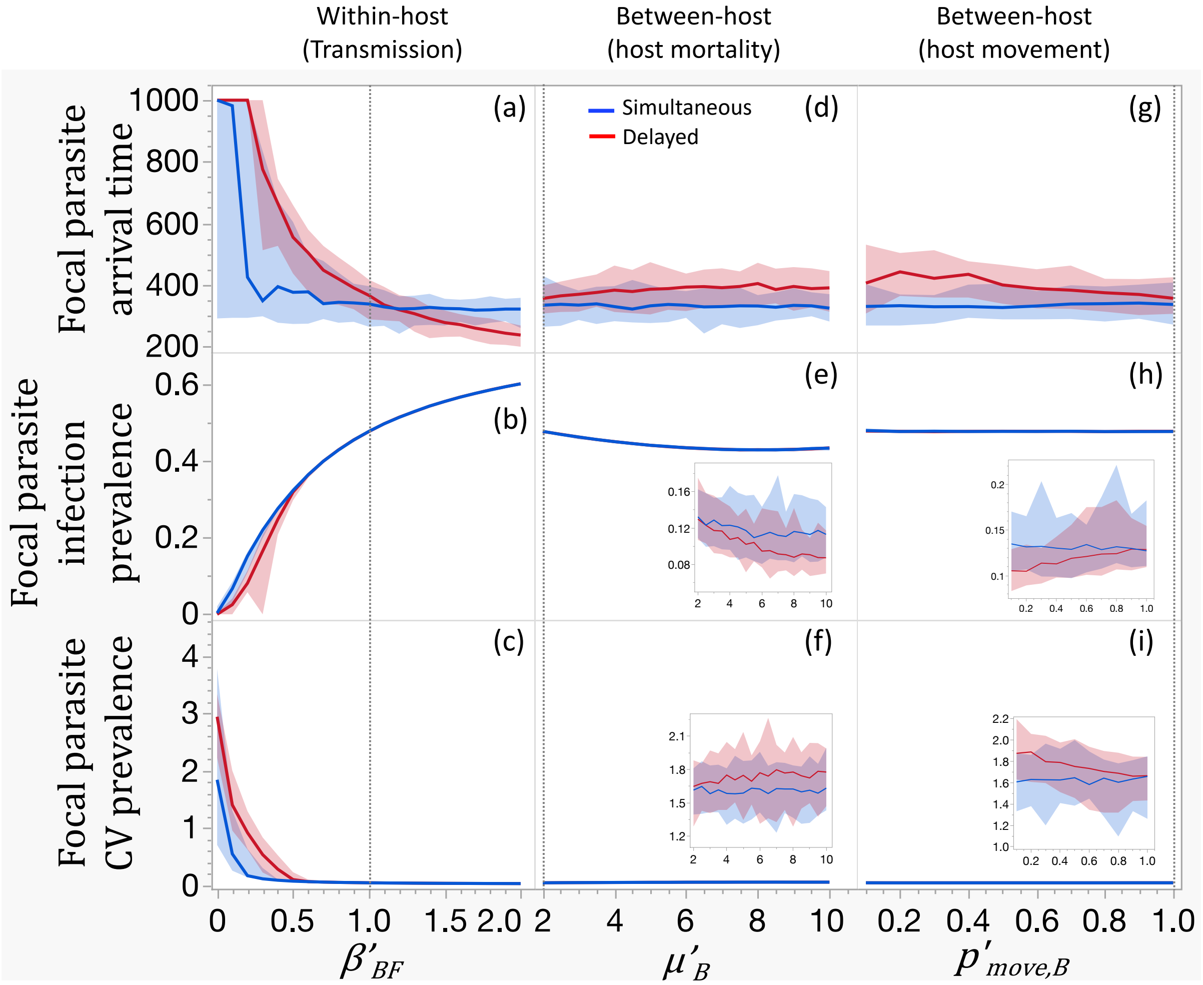

**Figure S2.** Equivalent to Figure 3 in the main paper, but assuming additive co-infected mortality (i.e., $\mu_C = \mu_F + \mu_B$). Model output, showing the effects of three types of parasite interactions by the background parasite on the spatial spread of the focal parasite at equilibrium, under two co-circulation scenarios (simultaneous introduction or delayed introduction of the focal). **(a)-(c)**: background parasite alters host susceptibility to the focal parasite via changes in $\beta'_{BF}$; **(d)-(f)**: background parasite increases host mortality via changes in $\mu'_B$; **(g)-(i)**: background parasite reduces host movement probability via changes in $p'_{move,B}$). Epidemiological metrics for the focal parasite relate to the time to spread from Patch 1 to the end patch of the network (top row), mean landscape-wide infection prevalence (middle row) and mean between-patch coefficient of variation of prevalence (standard deviation/mean; bottom row). Solid lines show the median values of 50 replicated simulations for each parameter value, shaded areas show the 95% quantiles of those values. The vertical dotted lines show the baseline (non-interaction) parameter values. Small insert graphs in panels e,f,h,i show zoomed-in versions of the results.

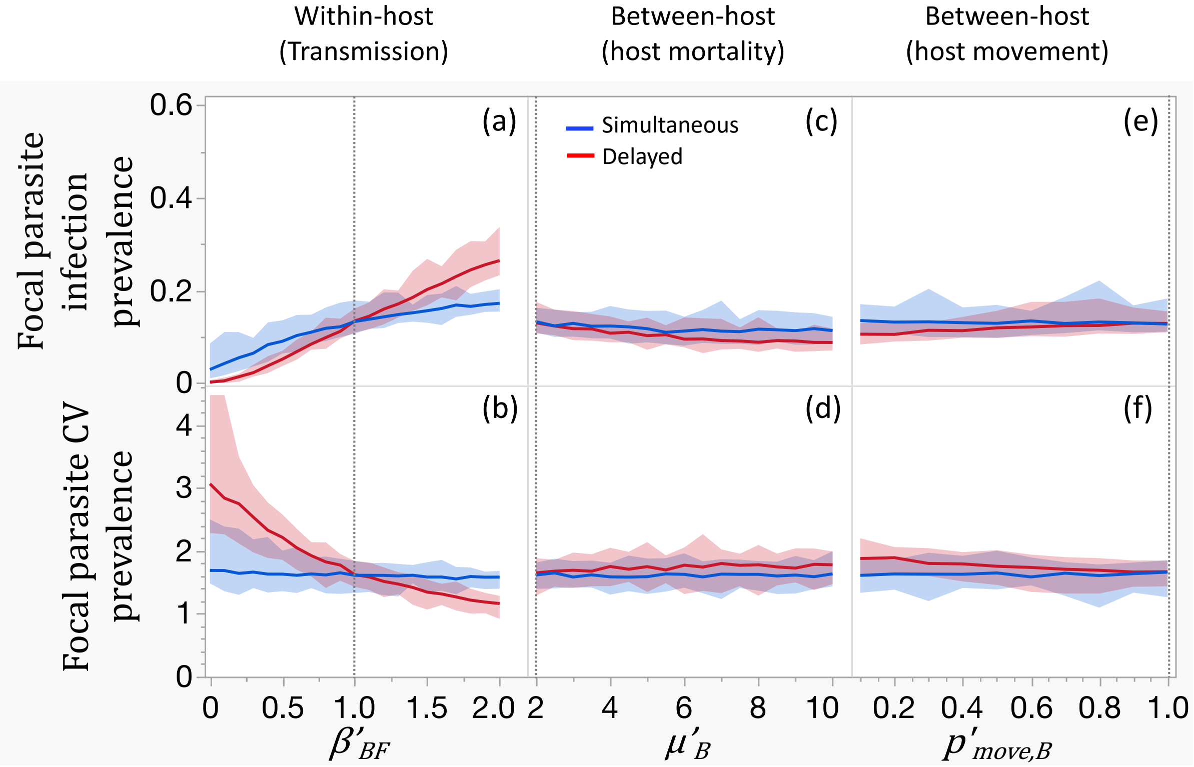

**Figure S3.** Equivalent to Figure 3 in the main paper, but for transient dynamics (sampled 100-120 time points after introduction of the focal parasite). Model output, showing the effects of three types of parasite interactions by the background parasite on the spatial spread of the focal parasite at equilibrium, under two co-circulation scenarios (simultaneous introduction or delayed introduction of the focal). **(a)-(b)**: background parasite alters host susceptibility to the focal parasite via changes in $\beta'_{BF}$; **(c)-(d)**: background parasite increases host mortality via changes in $\mu'_{B}$; **(e)-(f)**: background parasite reduces host movement probability via changes in $p'_{move,B}$). Epidemiological metrics for the focal parasite relate to mean landscape-wide infection prevalence (top row) and mean between-patch coefficient of variation of prevalence (standard deviation/mean; bottom row). Data on the time taken for the focal parasite to spread to the end patch of the network are not shown (equivalent to the top row in Figure 3), as the focal parasite did not arrive in the end patch during the transient phase. Solid lines show the median values of 50 replicated simulations for each parameter value, shaded areas show the 95% quantiles of those values. The vertical dotted lines show the baseline (non-interaction) parameter values.

### 2) Experimental details

#### *2.1 Biological system*

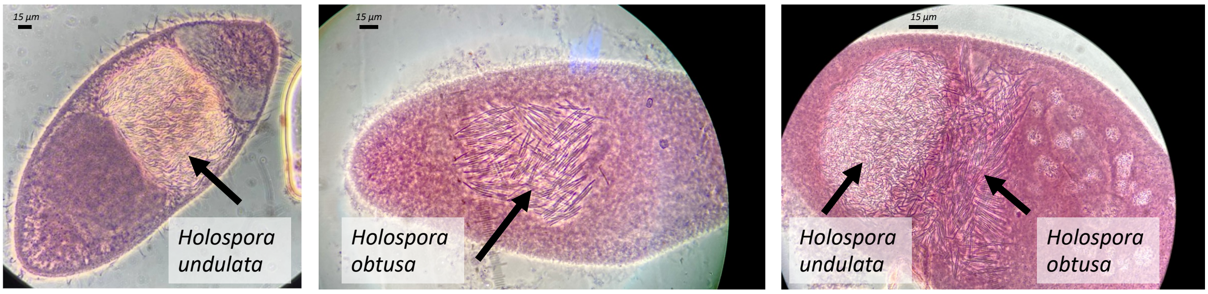

**Figure S4**. Single *Paramecium caudatum* individuals, with fully established *Holospora* infections. **(a)** Single *H. undulata* infection in heavily inflated micronucleus (MIC). **(b)** Single *H. obtusa* infection in macronculeus (MAC). **(c)** Co-infection with the two parasites in MIC and MAC, respectively. In all images, hosts carry large numbers (>>100) of elongated infectious forms of the parasites (10-15µm), S-shaped for *H. undulata* and rod-shaped for *H. obtusa*.

#### *2.2 Co-inoculation test: Impact of the background parasite on focal parasite transmission*

We tested if the presence of our background parasite *H. obtusa* affected the infection success of the focal parasite *H. undulata*. Inocula of *H. obtusa* and *H. undulata* were prepared by crushing >500 ml of infected stock culture of each parasite. After centrifugation (3.000G for 20 min), pellets containing infected cells were resuspended and subjected to mechanical lysis by agitation for 1 min 45 seconds in a mixer mill. For each resulting inoculum, we determined the concentration of infectious forms by counting the number of bacterial cells in a hemocytometer. We then adjusted the two inocula to the same concentration (by adding sterile Volvic™ water).

Inocula were added to 2 ml of uninfected *Paramecium* (c. 400 individuals, *P. caudatum* strain C084) in a 50 ml tube. All replicates received *H. undulata* inoculum, at four concentrations (25%, 50%, 75% and 100%), with 100% corresponding to 2.6 x $10^5$ infectious forms. We simultaneously added *H. obtusa* inoculum at four adjusted concentrations (0%, 25%, 50%, 75%), in the following 10 *H. undulata* (%) - *H. obtusa* (%) combinations: 25-0, 25-25, 25-50, 25-75; 50-0, 50-25, 50-50; 75-0, 75-25; 100-0.

Each combination was replicated 3 times. Infection success of *H. undulata* was determined 7 days post-inoculation, by fixing samples of c 30 individuals from each replicate, and checking their infection status under the microscope. Variation in the proportion of infected hosts was analysed using a LMM (binomial error and logit link, and correcting for overdispersion), with *H. undulata* inoculum concentration as explanatory factor and *H. obtusa* concentration as covariate.

We find that increasing *H. obtusa* dose caused decreasing infection probability of *H. undulata* ($\chi^2$ = 20.2, df = 1, p < 0.0001), from c. 60% in single inoculation to c. 40% at the maximum *H. obtusa* dose (Figure S5). Thus, the presence of *H. obtusa* transmission stages in the inoculum reduced infections of the focal parasite *H. undulata*. There was no significant effect of *H. undulata* dose ($\chi^2 = 4.2$, df = 1, $p > 0.2$) on infection probability. Very similar results were obtained, when *H. undulata* dose was fitted as a covariate rather than a factor (not shown). Note that two replicates were lost during handling. Data on the reciprocal effect of the focal parasite on the background parasite's transmission (J. Zabalegui, unpubl. data) will be published elsewhere.

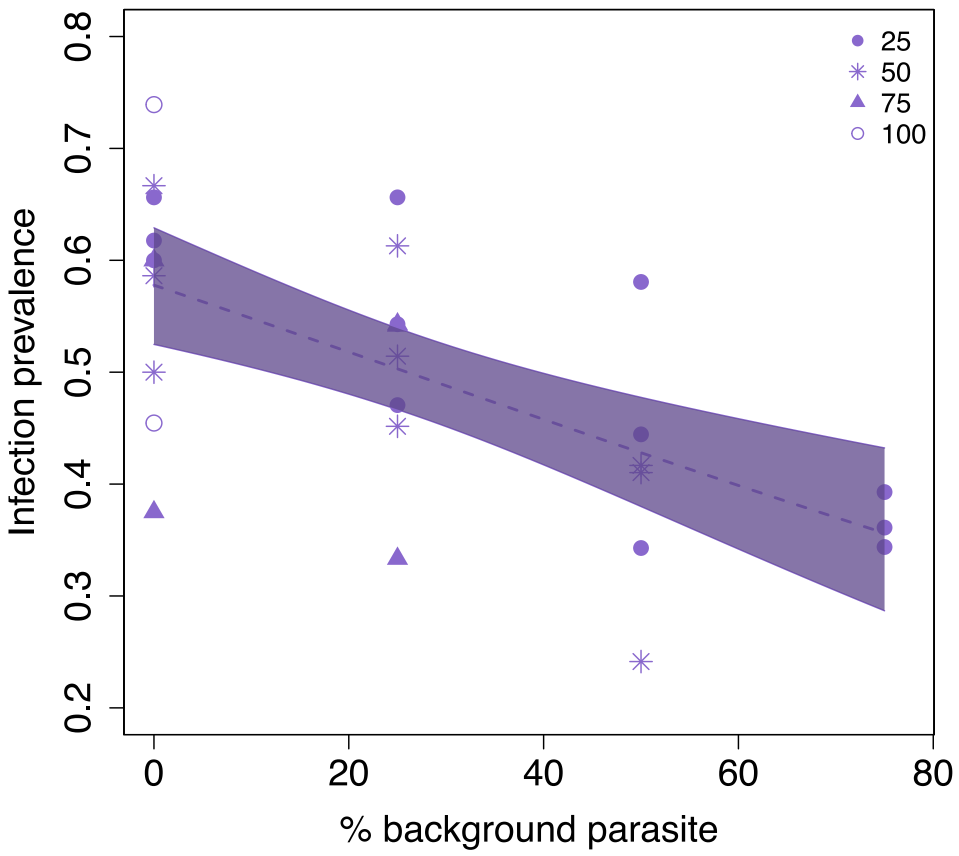

**Figure S5.** Infection success of *H. undulata* as a function of the dose of *H. obtusa*, simultaneously added to the *H. undulata* inoculum. The regression line is taken from a GLM (simplified model, with only *H. obtusa* dose as explanatory variable). The different symbols refer to four different *H. undulata* doses (25-100%).

#### *2.3 Landscape experiment*

#### *2.3.1 Spatial spread of uninfected host vs parasite*

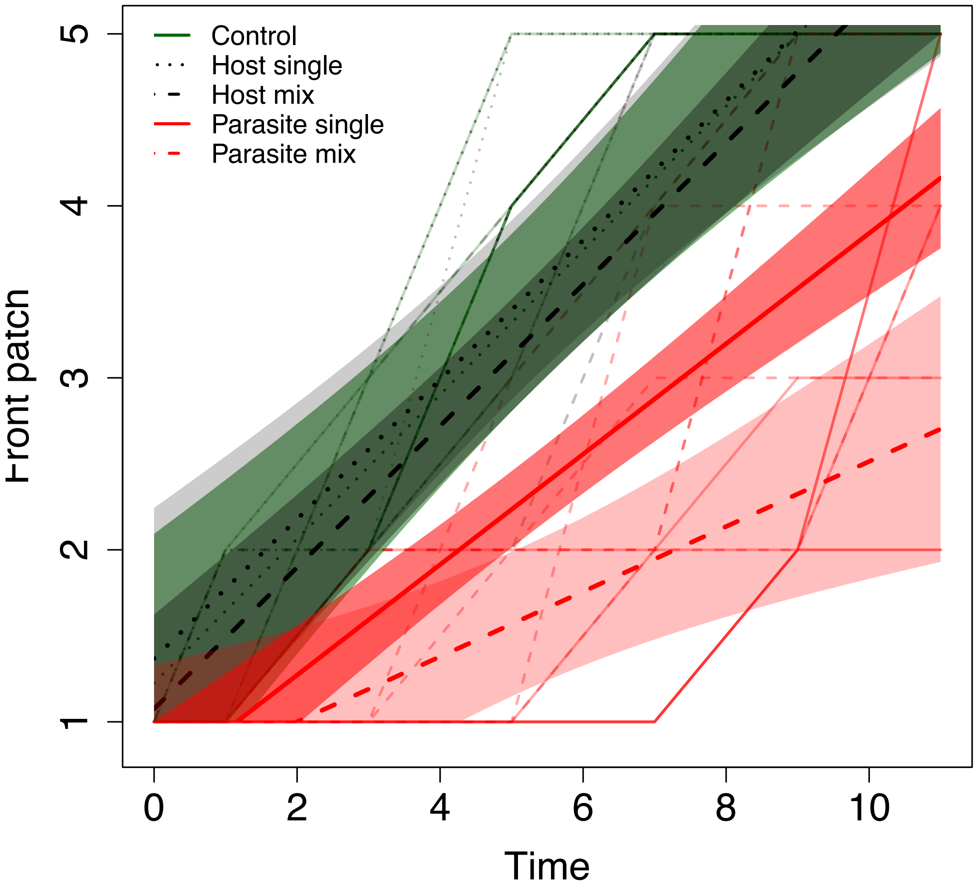

**Figure S6**. Rates of spatial spread of uninfected host (*P. caudatum*; green lines) and the focal parasite (*H. undulata*; red lines) into 5-patch landscapes. The front position is plotted against time (= number of 3h opening events), and thus the slope of the relationship describes the rate of spread. The spread of uninfected hosts is shown for uninfected control landscapes and for landscapes with the parasite (single- or mixed-parasite treatment). The spread of the parasite is shown for the two parasite treatments. The graph shows trajectories for each replicate landscape (thin lines), as well as the mean trajectories for the 3 host and 2 parasite groups (thick lines).

#### *2.3.2 Spatiotemporal dynamics of the spread of infection of the focal parasite (H. undulata)*

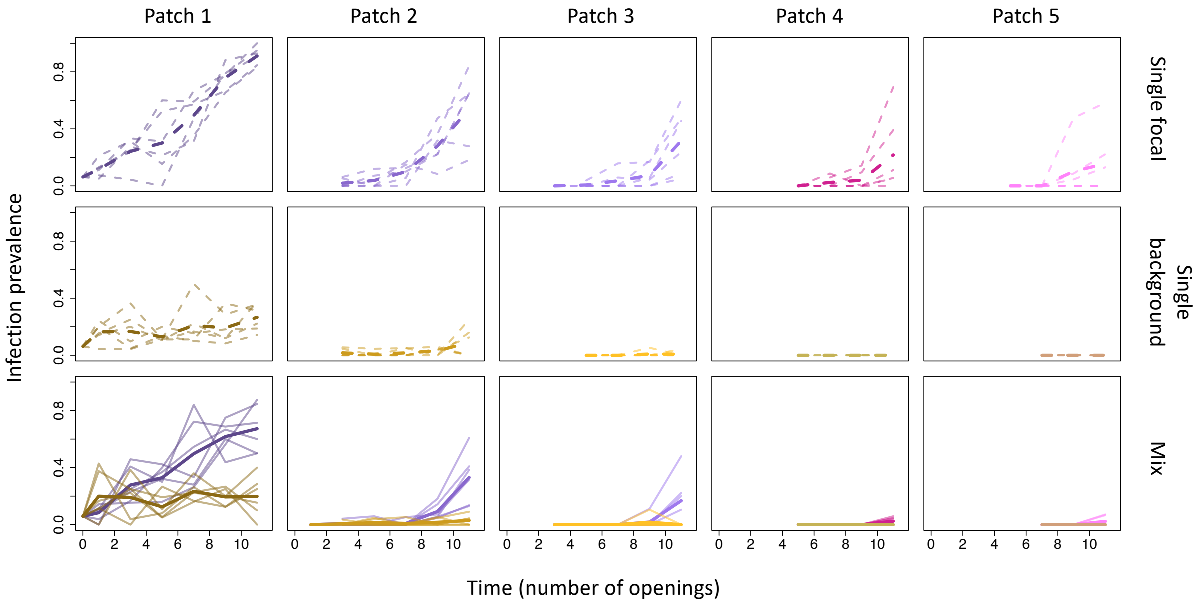

**Figure S7.** Spatiotemporal dynamics of the spread of infection in the single focal parasite treatment (*H. undulata*, top row), the single background parasite treatment (*H. obtusa*, middle row) and the mixed-parasite treatment (*H. undulata* (purple) and *H. obtusa* (yellow); bottom row). The plots show the trajectories of infection prevalence over time (= three weekly 3h opening events over the course of 4 weeks) in the 5 patches in each landscape. Infected hosts were introduced into the first patch, and the parasite then naturally spread into the downstream patches, travelling together with dispersing infected hosts. In each panel, the thin lines show the trajectories for individual replicates, the thick lines represent the mean trajectory.

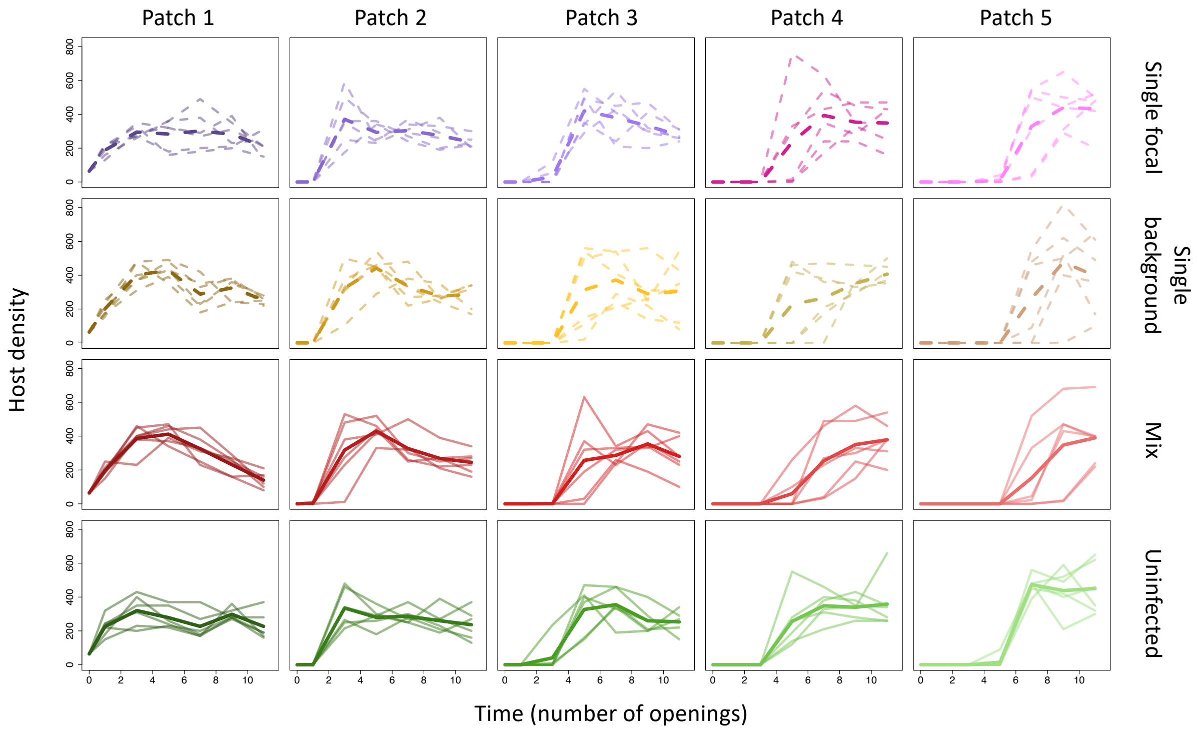

**Figure S8.** Spatiotemporal dynamics of host density (infected and uninfected combined) in the single focal parasite treatment (*H. undulata*, top row), the single background parasite treatment (*H. obtusa*, second row), the mixed-parasite treatment (*H. undulata (*purple) and *H. obtusa* (yellow); third row), and in the uninfected control treatment (bottom row). The plots show the trajectories of density over time (= three weekly 3h opening events over the course of 4 weeks) in the 5 patches in each landscape. Infected and uninfected hosts were introduced into the first patch; downstream patches were then colonised by dispersing hosts. In each panel, the thin lines show the trajectories for individual replicates, the thick lines represent the mean trajectory.

**Table S3** Linear Mixed Model analysing variation in spatial spread rate of the focal parasite (*H. undulata*). Variation in parasite front position was analysed as a function of time and parasite treatment (single- vs mixed-parasite). For random factors, variance (Var) and standard error (SE) are provided. 'Time' is taken as the number of times connections were opened between the patches in a landscape, allowing the dispersal of *Paramecium*.

| **Advancing-front position** | | | |
|---|---|---|---|
| *Random effects:* | Var (± SE) | | |
| Landscape identity | 0.01 ± 0.11 | | |
| Block | 0.0001 ± 0.01 | | |
| *Fixed effects:* | d.f. | $\chi^2$ | p |
| Parasite treatment | 1 | 9.70 | 0.001 |
| Time | 1 | 186.6 | <0.001 |
| Parasite treatment * Time | 1 | 6.98 | 0.008 |

**Table S4** General Linear Mixed Model analysing variation in focal parasite infection prevalence (*H. undulata*) at the final time point, as a function of patch position (1 to 5) and parasite treatment (single- vs mixed-parasite). For random factors, variance (Var) and standard error (SE) are provided.

| **Infection prevalence** | | | |
|---|---|---|---|
| *Random effects:* | Var (± SE) | | |
| Landscape identity | 0.61 ± 0.78 | | |
| Block | 0.04 ± 0.21 | | |
| *Fixed effects:* | d.f. | $\chi^2$ | p |
| Parasite treatment | 1 | 7.28 | 0.006 |
| Patch position | 1 | 217.1 | <0.001 |
| Parasite treatment * Position | 1 | 1.53 | 0.21 |

#### *2.3.3 Background parasite dynamics (H. obtusa)*

Following the protocol described in the main text, we set up an additional 6 replicate landscapes, with the background parasite starting from Patch 1 in a single-parasite treatment (*H. obtusa*). Thus, comparison with the mixed-parasite treatment allows us to assess the impact of the focal parasite (*H. undulata*) on *H. obtusa*. For details on the type of analyses, see main text.

Just like the spread of the focal parasite being negatively affected by the background parasite (see main text), we find the reverse trend of *H. obtusa* being negatively affected by the focal parasite (*H. undulata*), although less pronounced. Thus, while *H. obtusa*'s landscape-wide infection prevalence was generally lower in mixed (4.6 ± 1.8% SE) than single-parasite treatments at the end of the experiment (7.2 ± 2.2% SE; Figure S9b, Table S6), its spatial spread remained relatively unaffected by the focal parasite (Figure S9a). This is mainly because *H. obtusa* on its own (single-parasite treatment) did not succeed in spreading very far into the landscapes (see also Figure S7). As a consequence, the additional presence of the focal parasite (*H. undulata*) had had no significant effect (Table S5).

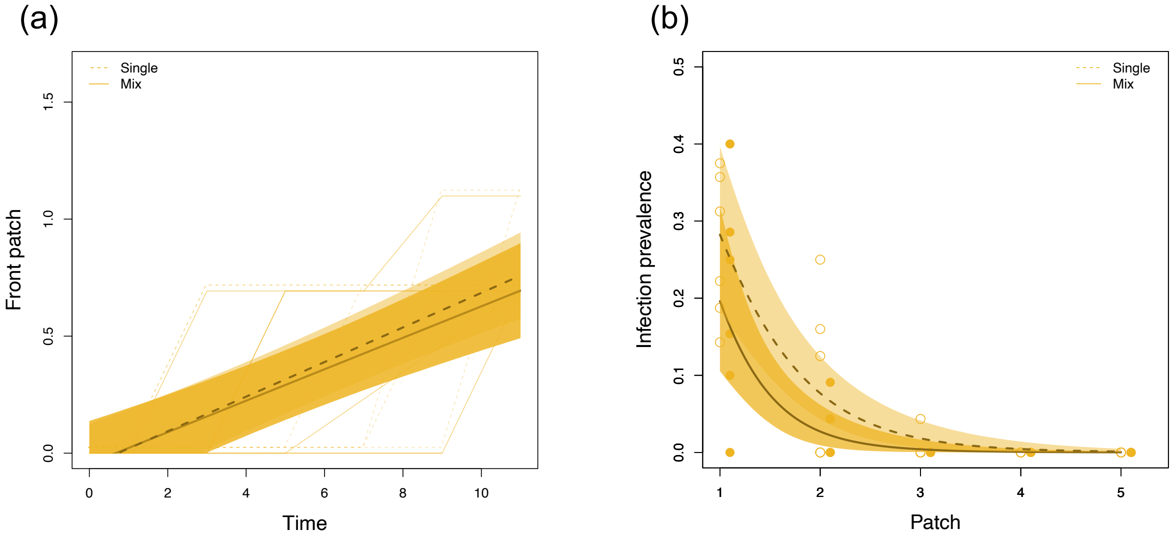

**Figure S9**. **(a)** Rate of spatial spread of the background parasite (*H. obtusa*) into the 5 patch landscapes, in single-parasite (dashed lines) and mixed-parasite (solid lines) treatments. The front position is plotted against time (= number of 3h opening events), and thus the slope of the relationship describes the rate of spread. **(b)** Infection prevalence of the background parasite (*H. obtusa*) across the 5 patches in single-parasite (dashed lines) and mixed-parasite (solid lines) treatments. The graphs show raw data (thin lines and points) as well as model predictions for the two treatments (median and 95% range).

**Table S5** Linear Mixed Model analysing variation in spatial spread rate of the background parasite (*Holospora obtusa*). Variation in parasite front position was analysed as a function of time and parasite treatment (single- vs mixed-parasite). For random factors, variance (Var) and standard error (SE) are provided. Block random factor was removed due to the negligible variance effect causing singular fit.

| **Advancing-front position** | | | |
|---|---|---|---|
| *Random effects:* | Var (± SE) | | |
| Landscape identity | 0.02 ± 0.15 | | |
| *Fixed effects:* | d.f. | $\chi^2$ | p |
| Parasite treatment | 1 | 0.05 | 0.82 |
| Time | 1 | 92.53 | <0.001 |
| Parasite treatment *Time | 1 | 0.20 | 0.65 |

**Table S6** General Linear Mixed Model analysing variation in background parasite infection prevalence (*Holospora obtusa*) across patches at the final time point. Variation in parasite front position was analysed as a function of patch position (1 to 5) and parasite treatment (single- vs mixed-parasite). For the random factor, variance (Var) and standard error (SE) are provided. Block random factor was removed from the full model due to the negligible variance effect causing singular fit.

| **Infection prevalence** | | | |
|---|---|---|---|
| *Random effects:* | Var (± SE) | | |
| Landscape identity | 0.17 ± 0.41 | | |
| *Fixed effects:* | d.f. | $\chi^2$ | p |
| Parasite treatment | 1 | 3.09 | 0.07 |
| Patch position | 1 | 52.48 | <0.001 |
| Parasite treatment * Position | 1 | 1.34 | 0.24 |

#### *2.4 Supplementary experiments*

The simulation model revealed possible epidemic-blocking spatial priority effects emerging when the focal parasite arrives in the landscape with a temporal delay. In two supplementary experiments reported here, we investigated the consequences of sequential arrival in our experimental model system.

#### *2.4.1 Experiment 1: Simultaneous vs delayed introduction of focal parasite in single microcosms*

In a first experiment, we simulated different scenarios of simultaneous and sequential arrival of our two parasites, using single microcosms (50 ml tubes). We established the following treatments, with the focal parasite (*H. undulata*) introduced either (i) alone, or together with background parasite (*H. obtusa*) either (ii) simultaneously, (iii) with a 1-week delay, or (iv) with a 2-week delay after the background parasite (Figure S10). Parasite introduction was performed by pipetting a given number of infected hosts into each microcosm, initially containing 200 uninfected hosts (in 1 ml of culture).

*Single-parasite*

| *B* | *F* |
|---|---|
| 0 | 5 |
| 0 | 25 |
| 0 | 45 |
| 0 | 50 |

*Mixed-parasite*

*Simultaneous*

| *B* | *F* |
|---|---|
| 45 | 5 |
| 25 | 25 |
| 5 | 45 |

*1-week delay*

| *B* | | *F* |
|---|---|---|
| 45 | | 5 |
| 25 | | 25 |
| 5 | | 45 |

*2-week delay*

| *B* | | *F* |
|---|---|---|
| 45 | | 5 |
| 25 | | 25 |
| 5 | | 45 |

**Figure S10.** Experimental setup in supplementary experiment 1, with single-parasite treatment (focal parasite *H. undulata*) and mixed parasite treatments (focal parasite and background parasite *H. obtusa*). In the mixed-parasite treatment, the focal parasite (*F*) was introduced simultaneously with the background parasite (*B*), or with a 1-week or 2-week delay. Different combinations of numbers of hosts (infected with either *B* or *F*) were tested.

To test effects of 'numerical contingency' of interaction effects, we further varied the (ratio of) the initial number of infected hosts pipetted into the microcosms, from 5 to 50 infected hosts (Figure S10). For the simultaneous treatments, three initial ratios of focal vs background numbers were established: 5-45, 25-25, 45-5. For the two delay treatments, we used these same ratios, but shifted in time. We hypothesised that the focal parasite would be at a particular disadvantage when in minority (5-45 treatment), or conversely, overcome a disadvantage when at initial numerical superiority (45-5 treatment). Note that we did not establish delayed single introductions of the focal parasite, due to logistic constraints. Over the course of the experiment, the initial 1-ml volume was progressively increased by adding fresh medium until the final volume of 30 ml was reached after 3 weeks.

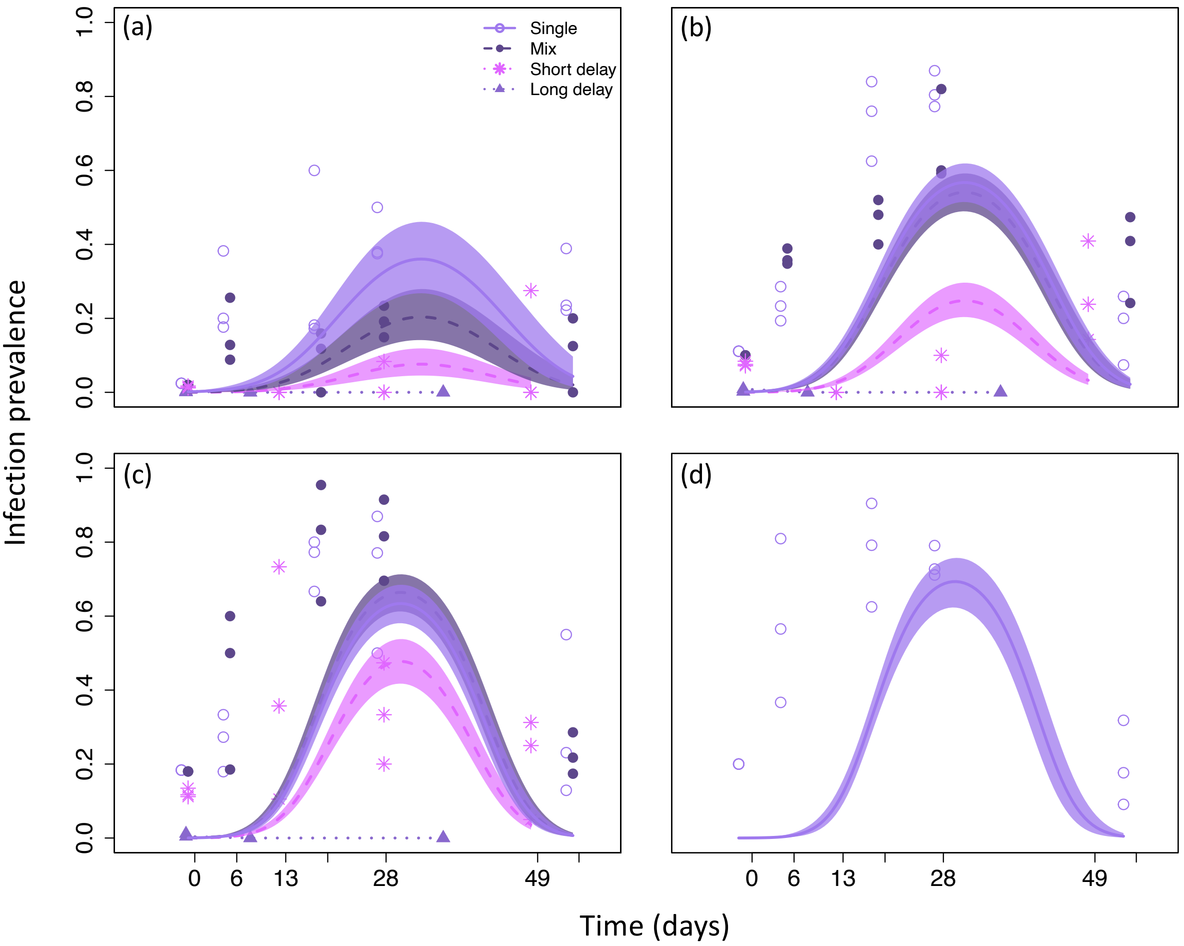

**Figure S11.** Time course of the spread of infection with the focal parasite *H. undulata* in single microcosms under contrasting epidemiological scenarios. The focal parasite was either introduced alone (single-parasite treatment) or together with the background parasite (*H. obtusa*), simultaneously or with a 1-week or 2-week delay (different curves in each panel). Each panel has different numbers of infected hosts introduced: **(a)** 5 focal + 45 background; **(b)** 25 focal + 25 background; **(c)** 45 focal + 5 background; **(d)** 50 focal. Statistical analyses (GLMM with binomial errors: time, time$^2$ and treatment as explanatory factors, replicate ID as a random factor) were carried out separately for each of panels (a) to (c). Treatment effects were significant in all 3 analyses ($\chi^2 > 35.3$, df = 2, p < 0.0001; the 2-week delay treatment was not included in analysis, as it produced consistent failure of focal parasite spread). In (d), only time and time$^2$ were fitted as the background parasite was not present. In all panels we show model predictions with median trajectory and 95% range.

We found that the initial presence of the background parasite caused a significant reduction of focal parasite prevalence (Figure S11; see figure legend for statistical analyses). When introduced simultaneously (red), focal infection outbreak trajectories were similar to that in single (=control) treatment (blue), with average peak prevalence being slightly higher in single-parasite (71.5 ± 5.1% SE) than that in the mixed treatment (56.6 ± 10% SE). When introduced with a delay, we found indications of a strong priority effect of the background parasite. With a 1-week delay, the focal parasite remained at relatively low infection prevalence (≤ 20%); following a 2-week delay, the focal parasite completely failed to establish. However, we cannot entirely rule out the possibility that the increase in host resistance against the focal parasite was caused by other changes in the host population, unrelated to the presence of the background parasite. This could be explicitly tested in future by introducing the focal parasite in background-parasite free populations after 1 and 2 weeks, respectively.

We further found a clear signature of the size of the initial cohort of infected hosts arriving in the host populations. Namely, when only 5 hosts infected with the focal parasite were introduced, subsequent epidemics peaked already at values of c. 30% infection on average, compared to >60% when 25, 45 or 50 individuals were introduced.

##### ***2.4.2 Experiment 2: Spatial priority effects in mini-landscapes***

We investigated possible spatial priority effects in 2-patch mini-landscapes, under natural dispersal. Under this scenario, the focal parasite (*H. undulata*) and background parasite (*H. obtusa*) were placed in opposite tubes, with different numbers of infected and uninfected hosts mixed at the beginning of the experiment. Thus, unlike in supplementary experiment 1 above, any delay in focal parasite arrival into a patch was naturally determined by the dispersal of infected hosts.

We set up 12 landscapes, with initially 2000 individuals (in 30ml) in each patch. In one set of 6 landscapes, the prevalence of the focal parasite in its source patch was set to 45%, while background parasite frequency in the neighbour patch varied from 0% to 5%, 15% and 45%. In a second set of 6 landscapes, focal parasite frequency in its source patch varied from 5% to 5%, 15% and 45%, while the background parasite prevalence in the neighbour patch was set to 45%. For each combination, we set up 2 replicate landscapes. Over the course of 3 weeks, connections were opened 3 times per week, for 3h. Infection prevalences were tracked, as described above and in the main text. We then analysed the performance of the focal parasite in the neighbour patch, as a function of its own initial prevalence in the source patch and the initial prevalence of the background parasite in the neighbour patch.

Generally, we detected the focal parasite in the neighbour patch during the first week (i.e., after 1-3 openings of the connection). As in supplementary experiment 1, the presence of the background parasite tended to limit focal parasite outbreaks (Figure S12). There was a significant effect of initial background-parasite frequency in the neighbour patch on focal parasite infection levels in that patch (background parasite level: $\chi^2$ = 15.9, df = 3, p = 0.0012). Essentially, focal parasite spread was increasingly reduced with increasing initial frequencies of the background parasite, indicative of a spatial priority effect. Thus, average maximum prevalences of c. 40-45% in the absence or with low initial frequency of the background parasite dropped to values as low as 12.5% when the background parasite was initially frequent (mean: 31%).

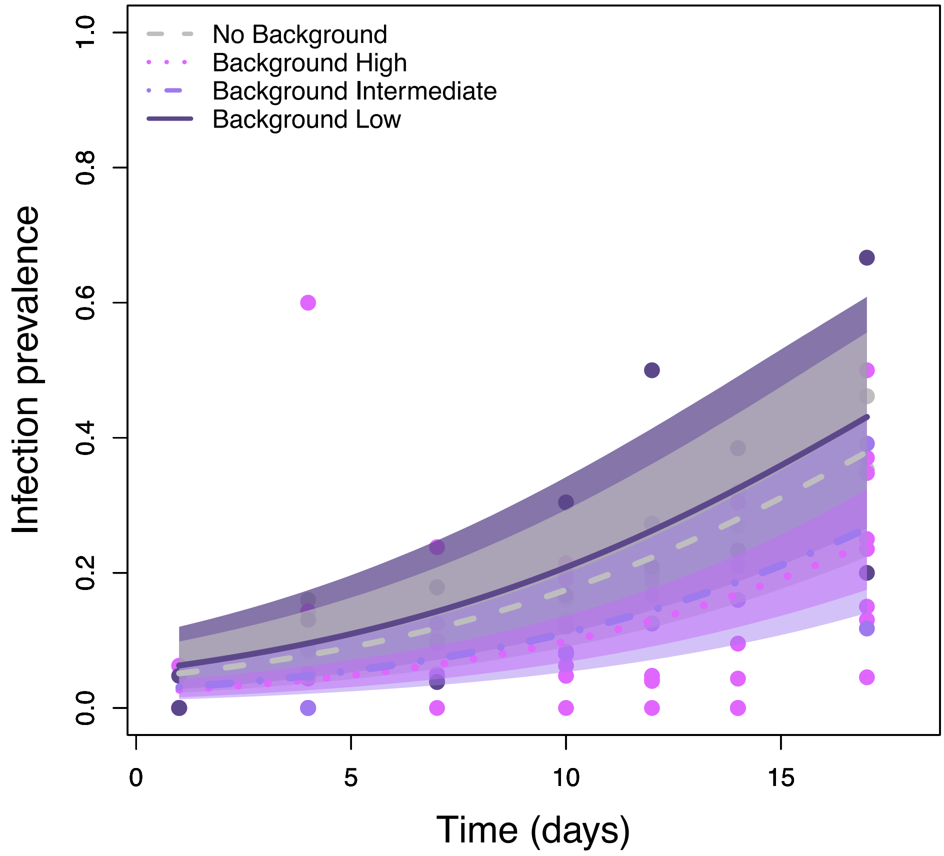

**Figure S12.** Time course of infection levels of the focal parasite *H. undulata* in Patch B, when being naturally introduced from Patch A via dispersing infected hosts. Patch B initially harboured different frequencies of the background parasite (*H. obtusa*), ranging from 0% to 5% ('low'). 15% ('intermediate') and 45% ('high'). Model predictions (median trajectory and 95% range) are shown for each of these initial conditions.
